# *In vivo* tractography of human neonatal white-matter pathways underlying hypothalamic and reward functions to study predispositions to neurodevelopmental conditions and obesity

**DOI:** 10.1101/2025.02.28.640038

**Authors:** Julie Nihouarn Sigurdardottir, J-Donald Tournier, Dafnis Batalle, Alexandra F. Bonthrone, Daan Christiaens, Jana Hutter, Lucillio Cordero-Grande, Andrew Chew, Chiara Nosarti, Jo Hajnal, Grainne McAlonan, Lucilla Poston, Mary Rutherford

## Abstract

White matter (WM) tracts of the reward, limbic, and autonomic systems implicate the hypothalamus, nucleus accumbens, ventral tegmental area and the amygdala and are associated with autism, ADHD, addiction and obesity. However, since most of these structures remain uncharacterised *in vivo* in human neonates, research on the early-life predispositions to these long-term “mind and body” conditions and the impact of common fetal exposures such as maternal obesity remains limited. Through the developing human connectome project, we acquired 3T brain diffusion and structural magnetic resonance imaging from healthy neonates born at-term to 137 normal-weight women (controls) and to 28 obese women and scanned at mean 40 weeks+6 days (+/-9 days) postmenstrual age (PMA). We first developed novel tractography protocols to reconstruct anatomical WM pathways for the neonatal medial forebrain bundle, ventral amygdalofugal pathway, amygdalo-accumbens fasciculus, stria terminalis and autonomic dorsal longitudinal fasciculus (DLF). We then quantified WM structure from the mean tract fibre bundle density (FD) and fibre cross-section (FC) and using regression path models evaluated WM change across PMA and the effects of antenatal obesity exposure and neonatal covariates. Lastly, we explored if neonatal WM FD and obesity exposure predicted child psycho-cognitive outcomes and anthropometry at 18 months. We show successful *in vivo* tractography of tracts with high topographical correspondence to adult histology, including in subcompartments of the hypothalamus and amygdala. The obesity exposure*PMA interaction was significant for mean FD in the bilateral amygdalo-accumbens fasciculus and right uncinate fasciculus. Males had larger FC in these same tracts bilaterally. Antenatal obesity exposure predicted lower cognitive scores and higher WHO weight and height z-scores at 18 months. Toddler reward-seeking temperament was correlated with higher weight zscore and was predicted by higher neonatal FD of the amygdalo-accumbens and uncinate fasciculi. Denser neonatal DLF predicted higher language and cognitive scores and fewer autistic traits at 18 months. In conclusion, we inform on neuroanatomical growth *in vivo* of discrete multisystemic regulatory networks and present evidence for early-life predispositions to psychological outcomes and obesity.

## Introduction

Brain areas and networks that contribute to weight gain through feeding behaviour and energy regulatory pathways have also have been directly or indirectly implicated in executive function, impulse control and inhibition, social cognition and behaviours and more broadly in the phenotypes of autism and attention-deficit hyperactivity disorder (ADHD; Hughes and Ensor, 2007). While their interaction with environmental factors is assumed to be complex, such neurological overlap could explain the reported incidence of comorbidity in these “mind and body” outcomes among youth and adults (Kahathuduwa et al., 2019; Garcia-Argibay et al., 2022, 2024). However, to date there is little evidence which offers the underpinning in the human brain of an early-life predisposition to both health domains and their potential for comorbidity. Here, capitalising on neonatal diffusion neuroimaging and tractography *in vivo*, we propose a study to reveal this across a comprehensive set of interlinked brain pathways and their association with outcomes in infancy. We also translate it into an exploration of the effect of maternal obesity, a rising fetal exposure associated with these offspring phenotypes.

A primary candidate for such neurological underpinning is the mesolimbic/reward network. It encompasses the ventral tegmental area (VTA), nucleus accumbens (NAcc) and the amygdala and receives regulatory control from the prefrontal cortex. The amygdala is implicated in the stress response and in pleasure-inducing behaviour, including eating palatable food (Ulrich-Lai et al., 2010). The NAcc and reward networks were previously shown to have an aberrant structural connectivity between the amygdala with other areas of the reward system (mPFC) in pleasure-inducing contexts, as studied in models of chronic early life adversity (Bolton et al., 2018). Together, this network is predominantly implicated in reward-seeking behaviour and in integrating social reinforcement learning through dopamineric activity (Solié et al., 2021). For example, it modulates addictions to food, drugs and sex (Wang et al., 2005; Kenny, 2011; Volkow et al., 2017), contributes to anhedonia following stress (Russo and Nestler, 2013; Friedman et al., 2014), in the social aspects of autism (Bariselli et al., 2018; Supekar et al., 2018) and is altered in adults with ADHD (Volkow et al., 2011). It has also been shown to explain the association between the genetic liability for Body Mass Index (BMI) and higher odds for ADHD (Martins-Silva et al., 2021). Two WM tracts of this network are the medial forebrain bundle (MFB), which links the VTA and NAcc, and the amygdalo-accumbens fasciculus (Am-NAcc) which links the amygdala to the NAcc (Rigoard et al., 2011; Supekar et al., 2018).

The hypothalamus is formed of multiple nuclei and also plays a key role in neurodevelopmental and metabolic phenotypes. From birth, growth and the stress response are regulated via its interaction with the pituitary and the adrenal glands. It relays signals within the reward network with the aformentioned MFB traversing the lateral hypothalamic compartments. Within the limbic system, the hypothalamus links directly to the amygdala via the long loop of the Stria Terminalis (ST) and via the shorter fiber bundle of the ventral amygdofugal pathway (vAMFP), which have been implicated in motivated behavioural responses and feeding behaviour (Rollins et al., 2006; Choi et al., 2010; Krüger et al., 2015; Kamali et al., 2016; Gorka et al., 2018; Bech et al., 2020). The hypothalamus has also the pivotal function of regulating satiety, cardiovascular function, alertness and overall physiological homeostasis as the master regulator of the autonomic nervous system (ANS). For example, the vagal gut-brain pathway has received increased interest in research on food addiction (Gupta et al., 2020), metabolic/cardiovascular health (Bouret, 2012), and neurodevelopment including in the paradigm of antenatal susceptibility to psychopathology through the microbiome (Cryan and Dinan, 2012; Codagnone et al., 2019). One WM bundle of interest to this study is the dorsal longitudinal fasciculus (DLF) which contributes to the transmission between the hypothalamus and the vagus nerve (Halász, 2004; Rea, 2015).

The cingulum and uncinate fasciculus (UF) are other limbic WM fiber bundles included in this study of predispositions to obesity and psychological phenotypes. They are already defined using neonatal neuroimaging and tractography and they directly connect inhibitory orbitofrontal areas widely across other lobes. Both have been associated with reward processing in adults with ADHD who show lower activation in decisionmaking tasks (Yang et al., 2019). The UF is implicated in autism and conduct disorders of impulsivity and cognitive control (Olson et al., 2015) and with functional connectivity of the NAcc and reward sensitivity (Camara et al., 2010; Bjornebekk et al., 2012).

An anatomical and structural characterisation of the neonatal networks described above has not only the potential of improving our understanding of early-life origins of population psychological and physiological health but also study how specific antenatal environments shape their developmental trajectories. Currently, the rates of obesity in the female population of child-bearing age are reaching 35% globally (Chen et al., 2018; Kent et al., 2024). Converging evidence suggests an interaction between *in utero* exposure to maternal obesity and higher likelihood of offspring neurodevelopmental conditions and metabolic disorders, all of which are increasing world-wide (Li et al., 2016; Lei et al., 2019; Kong et al., 2020; Duko et al., 2024). Animal models of high-fat-diet induced obesity as well as human clinical evidence suggest the offspring have increased risk of obesity, metabolic and cardiovascular disease (Steculorum and Bouret, 2011; Bouret, 2012; Yu et al., 2013; Vogt et al., 2014; Park et al., 2020) but also of two-fold higher likelihood of autistic spectrum condition (ASC) and ADHD (Adane et al., 2016; Li et al., 2016). Children diagnosed with ASC or ADHD are also more likely to be overweight or obese than undiagnosed children (Broder-Fingert et al., 2014; Cortese et al., 2016) although this association is not always consistent (Dubnov-Raz et al., 2011) and/or may show sexual dimorphic effects (Barnhill et al., 2017).

Nevertheless, testing the hypothesis that early predispositions to neurodevelopmental and metabolic/obesogenic outcomes are increased following obesity exposure compared to normal-weight exposure has not been feasible due to the lack of methods for tracing,*in vivo*, the brain pathways described above and quantifying their structural properties when the influence of the post-natal environment is minimal. The hypothalamus, NAcc and VTA and these WM pathways have been understudied structures in the neonatal human population due to past limitations in resolution and sensitivity of structural and diffusion neuroimaging in the newborn. Additionally, neuroimaging in the neonate offers the advantage of excluding early childhood exposure confounding effects of illness, lifestyles linked to an obesogenic environment (e.g. parental feeding, low child exercise, picky eating) (Matheson et al., 2015), parenting strategies and parent mental health which could all be implicated in the causal pathway of interest (Dufford et al., 2021).

Here, we relied on state-of-the-art neonatal data collected from the developing human connectome project (dHCP) with the first aim to develop novel *in vivo* tractography protocols for the neonatal Am-NAcc, DLF, MFB, ST and vAMFP. The second aim was to assess the anatomical topography and the structural development of fiber bundles in the healthy term-born neonates using fixel-based analysis rather than conventional diffusion tensor-based metrics. This approach was selected to overcome biases in axonal micro and macrostructure quantification due to crossing and “kissing” fibres within the same voxel (Tournier et al., 2007; Jeurissen et al., 2014; Raffelt et al., 2017a), characteristic of the diencephalic WM. To improve statistical robustness of estimators we used statistical path regression modelling and studied the effects of sex, postmenstrual age, postnatal age, brain volume and birthweight centile on WM development in normal-weight exposed participants. Finally, in an attempt to understand antenatal effects of maternal obesity, we compared WM structure between exposed and non-exposed neonates and their relationship with child anthropometric and psychological outcomes at 18 months.

## Methods

### Participants

Parents of neonates provided written informed consent for the child to participate in the developing Human Connectome Project (dHCP; ethics approval: 14/LO/1169). Demographic data were collected at enrolment. Pregnancy and infant outcomes were retrieved from the medical records, discharge notes and questionnaires (Edwards et al., 2022) and a birthweight centile was calculated using INTERGROWTH-21st (v.1.3.5, Villar et al., 2014). The neonates included were considered healthy and from uncomplicated pregnancies, minimising other confounding antenatal exposures and adverse birth outcomes: From 962 participants in the dHCP database at the time of enquiry, we excluded non-singleton pregnancies, neonates of women with missing obstetric outcomes or BMI, who reported obstetric cholestasis, preeclampsia, Haemolysis, Elevated Liver enzymes and Low Platelets (HELLP), pregnancy induced hypertension and gestational diabetes mellitus. Also excluded were neonates born outside the range 37-42 weeks gestational age,with apgar < 7 at 5min (or missing) and who were admitted to intensive care. We only included those born to normal-weight (18.5 to 25 kg/m^2^) or obese women (>=30 kg/m^2^). Blinded to exposure group we reviewed the MRI quality and radiology reports and excluded poor quality data and neonates with germinal matrix hemorrhage, intraventricular hemorrhage and bilateral temporal horn cyst, >4 WM punctate lesions and congenital abnormalities (e.g. cardiac defect). The final sample count was 137 neonates born of normal-weight mothers and 28 of mothers with obesity. See Table 1 for maternal, birth and neonatal comparisons between the groups.

**Table 1:** Maternal and neonatal characteristics.

|  | Overall | Normal-weight (Controls) | Obese | p |
| --- | --- | --- | --- | --- |
| n | 165 | 137 | 28 |  |
| **Mother** |  |  |  |  |
| Mother's age (mean [SD]) | 33.12 (4.86) | 33.19 (5.03) | 32.79 (4.04) | 0.690 |
| Mother's ethnicity (%) |  |  |  | <0.001 |
| White | 101 (62.3) | 91 (67.9) | 10 (35.7) |  |
| Black | 22 (13.6) | 10 (7.5) | 12 (42.9) |  |
| Asian | 17 (10.5) | 16 (11.9) | 1 (3.6) |  |
| Chinese | 7 (4.3) | 7 (5.2) | 0 (0.0) |  |
| Mixed | 5 (3.1) | 3 (2.2) | 2 (7.1) |  |
| Other | 10 (6.2) | 7 (5.2) | 3 (10.7) |  |
| Mother's first language is English (%) | 86 (53.1) | 74 (55.2) | 12 (42.9) | 0.325 |
| Mother's age when last in FT education (median [IQR]) | 23.00 [21.00, 25.00] | 23.00 [21.00, 25.00] | 23.00 [21.50, 24.00] | 0.930 |
| Mother's BMI (median [IQR]) | 22.39 [20.50, 24.19] | 21.79 [20.20, 23.37] | 32.20 [30.86, 35.18] | <0.001 |
| Primiparous (%) | 91 (55.2) | 83 (60.6) | 8 (28.6) | 0.004 |
| **Delivery** |  |  |  |  |
| Meconium stained liquor (%) | 39 (23.9) | 32 (23.7) | 7 (25.0) | 1.000 |
| Delivery Mode (%) |  |  |  | 0.574 |
| Elective caesarian section | 20 (12.1) | 14 (10.2) | 6 (21.4) |  |
| Emergency caesarian section - in labour | 34 (20.6) | 29 (21.2) | 5 (17.9) |  |
| Emergency caesarian section - not in labour | 8 (4.8) | 6 (4.4) | 2 (7.1) |  |
| Instrumental delivery - Forceps | 30 (18.2) | 26 (19.0) | 4 (14.3) |  |
| Instrumental delivery - Ventouse | 12 (7.3) | 11 (8.0) | 1 (3.6) |  |
| Spontaneous vaginal delivery | 61 (37.0) | 51 (37.2) | 10 (35.7) |  |
| **Neonate** |  |  |  |  |
| Baby sex Female (%) | 70 (42.4) | 60 (43.8) | 10 (35.7) | 0.563 |
| GA at birth weeks (mean [SD]) | 40.19 (1.08) | 40.25 (1.05) | 39.87 (1.15) | 0.085 |
| Apgar 5 min (median [IQR]) | 10.00 [9.00, 10.00] | 10.00 [9.00, 10.00] | 10.00 [9.00, 10.00] | 0.550 |
| Baby head circumference (mean [SD]) | 34.49 (1.56) | 34.55 (1.54) | 34.19 (1.67) | 0.266 |
| Baby birthweight (kg) (mean [SD]) | 3.41 (0.48) | 3.40 (0.47) | 3.46 (0.52) | 0.522 |
| Birthweight centile (mean [SD]) | 53.33 (29.98) | 52.21 (30.07) | 58.81 (29.43) | 0.290 |
| PMA at scan (mean [SD]) | 40.83 (1.31) | 40.95 (1.31) | 40.22 (1.09) | 0.006 |

*Note:*
Mother's BMI (Body mass index) was booking BMI at ~ 12 weeks. Group-wise one-sided comparisons (t-test for parametric or Kruskal–Wallis for non-parametric continuous variables and Chi-squared/Fisher test for categorical variables) were performed to evaluate group-wise differences on maternal or delivery outcomes. Missing data: Maternal ethnicity=3, Maternal first language=3, Maternal education=16. Neonatal meconium=2. Birthweight centile was calculated from INTERGROWTH-21st standards. FT: Full-time, GA: gestational age, PMA: Postmenstrual age (e.g. GA at birth + chronological postnatal age)

Follow-up data was available for 129 of 137 infants at 18 months (see below).

### MRI of the newborn

All imaging data was acquired on a 3T Philips Achieva scanner within the Evelina Newborn Imaging Centre at St Thomas’ Hospital, London, and included structural (T1 and T2-weighted) and diffusion imaging (Edwards et al., 2022). The set up includes a bespoke neonatal imaging system (Hughes et al., 2017) which maximises signal-to-noise ratio by minimizing movement and incorporating a close fitting 32-channel receiver coil. Infants were scanned during natural sleep usually after a feed. Moulded silicone earplugs were placed in the infant ears in addition to neonatal earmuffs and an acoustic foam hood which was placed over the baby. A pediatrician or neonatal nurse was present throughout the scan and vitals of the infant monitored. T2-weighted images were reconstructed to a 0.5mm^3^ resolution (see Edwards et al., 2022). Diffusion data of 300 volumes were acquired during a 19 min scan through a Multi-shell High Angular Resolution Diffusion Imaging (HARDI) single-shot spin-echo echo-planar sequence (Hutter et al., 2018; Tournier et al., 2020) with multiband factor of 4, slice interleave factor 3, TE/TR:90/3800msec, Field of view 150 × 150 × 102 mm^3^, 1.5×1.5×3mm voxel resolution with 50% slice overlap, SENSE 1.2, Half-Fourier 0.85, 300 directions at b-value shells 0 (x20), 400(x64), 1000(x88) and 2600s/mm^2^(x128). Diffusion images were denoised and reconstructed to a 1.5mm^3^ isotropic resolution (Bastiani et al., 2019; Cordero-Grande et al., 2019), Gibbs ringing suppressed (Kellner et al., 2016) and motion and distortion corrected (Christiaens et al., 2021).

### Fixel-based Analysis (FBA)

For a detailed description of the pipeline and commands, see the Supplementary. In summary: The response functions for WM and CSF were estimated (Dhollander et al., 2019) using *MRtrix3* (Tournier et al., 2019). For each subject, a Fibre Orientation Distributions (FODs) map was obtained with 2-tissue multishell multi-tissue constrained spherical deconvolution (Jeurissen et al., 2014). The FODs were intensity normalised (Raffelt et al., 2017b) and a brain mask was also produced. FBA was implemented on the sample using the documented pipeline (Dhollander et al., 2021) and with minimal adaptation to the neonatal data. It consists of a template FOD generation, anatomical registration of regions of interest (ROIs) to the FOD template (Figure S1), fixel masks generation and thresholding (Figure S3) and the extraction of mean FD, logFC and FDC. The entire pipeline was performed in *MRtrix3* (version 3.0).

### Tracts of interest and Anatomical regions of inclusion and exclusion

We describe the tracts of interest as reported from the available literature at the onset of the study in the Supplementary. Tractography required mask regions of anatomical structures for seeding, exclusion and inclusion, both manually drawn or derived from atlas-based masks. The hypothalamus (with mamillary bodies), NAcc and VTA were manually segmented (JS), relying on previous adult MRI literature and atlases (Baroncini et al., 2012; Makris et al., 2013), see Figure 1 A. Other masks derived from the atlases adapted to the neonatal dHCP data were the amygdala, hippocampus, the anterior temporal lobe (medial and lateral), the mid and orbitofrontal WM and the thalamus (Makropoulos et al., 2014; Schuh et al., 2018). A study template T2 image was generated and the median of the subject atlas-based ROI masks from 10 randomly selected participants per group used to produce the FOD population template were registered to this T2 template. Other inclusion and exclusion “boxed” regions were manually drawn on the FOD template relying on the T2 template and a FOD-based directionally encoded color (DEC) map (Figure S2, *fod2dec*) (Dhollander et al., 2015).

**Figure 1:**
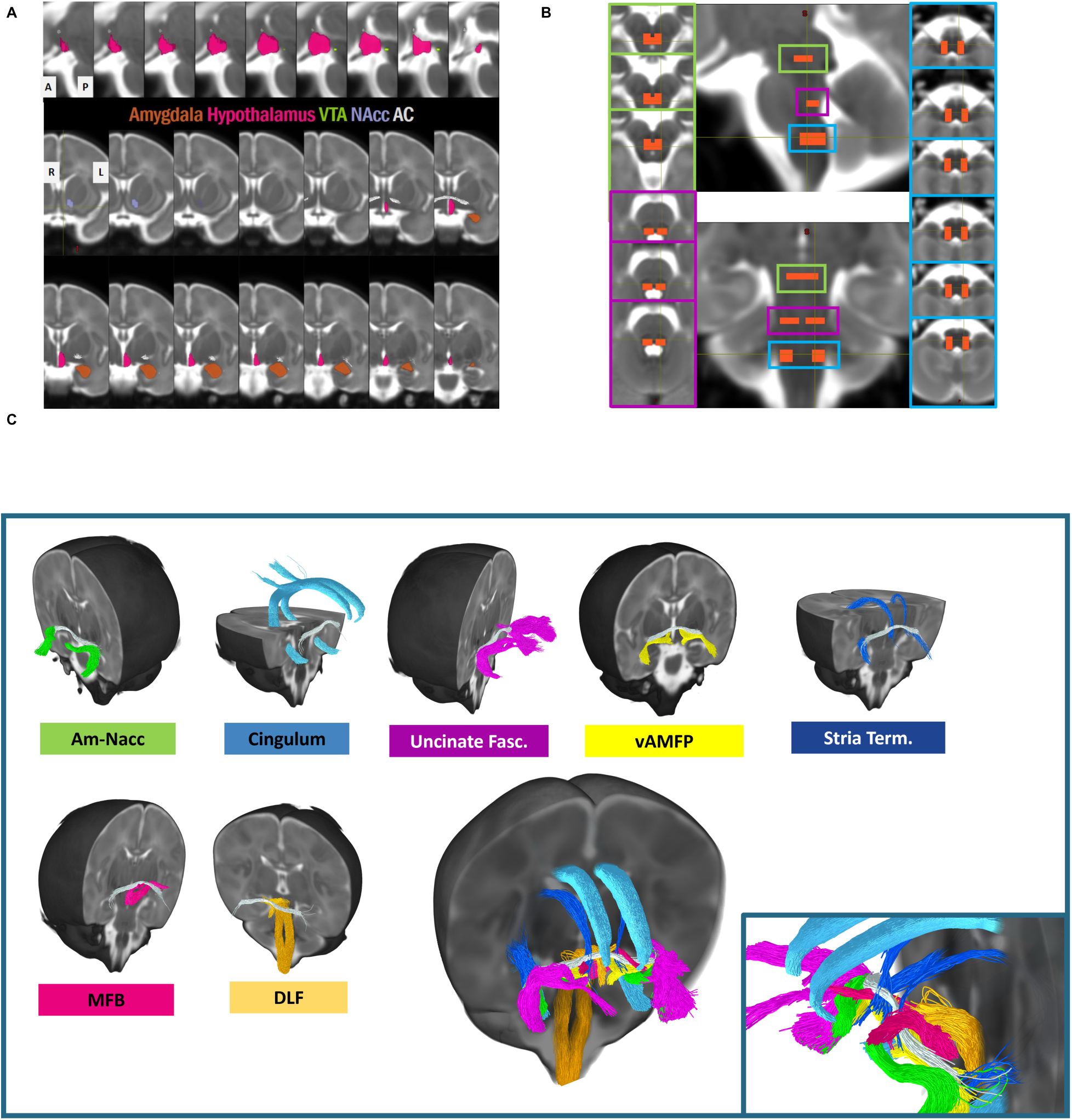
A. Manually drawn regions for tractography: hypothalamus (pink), ventral tegmental area (green, VTA), nucleus accumbens (lavender, NAcc) and the atlas-based segmentation of the amygdala (orange). Top row show sagittal planes through the hypothalamus in the lateral to medial direction. Bottom row are coronal slices arranged in the anterior to posterior directions. B. Regions of inclusion manually drawn for tractography of the Dorsal Longitudinal Fasciculus at the level of the midbrain periaqueductal grey matter (green), mid pons (fuchsia) and caudal medulla (blue) at a region assumed to include cranial nerve X (vagus) fibres, enlarged in Figure S4. Central top image is in sagittal view and bottom in coronal view. C: Tractography of fibre bundles shown overlaid on the population T2 template: Amygdalo-accumbens tract (Am-Nacc, green), cingulum (light blue), dorsal longitudinal fasciculus (DLF, orange), medial forebrain bundle (MFB,magenta), stria terminalis (Stria Term, dark blue), ventral amygdalofugal pathway (vAMFP, yellow) and uncinate fasciculus (fushia). Box: zoom view, excludes the left uncinate. The anterior commissure (grey) was generated in order to offer a reference tract for visualisation.

### Tractography protocols

All tracks were generated in template FOD space, using probabilistic tractography (iFOD2 algorithm) in *MRtrix3*. Protocols were adapted for each tract to minimise the number of spurious and false positive tracts and to account for the FOD derived signal being lower in regions where WM and GM mix within voxels in the subcortical areas studied here.

Since many tracts had not been delineated in the neonate previously and others have used WM atlases based on manual delineation rather than tractography, certain tracts of interest (DLF, MFB) were first generated in a sample processed adult brain from the Human Connectome Project. This helped determine the location of regions of inclusion at a resolution available in our cohort and we also relied on structural *ex-vivo* anatomical atlases of the fetus and neonate (Bayer and Altman, 2003). Anatomical correspondence of the “core” body of a streamline bundle within each tract was prioritised over the terminations of some fibre bundles which are known to “fan” out as they approach cortical surface (i.e. uncinate fasciculus) as this could influence the statistics when using mean FBA-derived FC/FD/FDC within fixel masks. A further step to minimise this was to utilise fixel thresholding (count of streamline per fixel) within the fixel masks for each tract. Unless otherwise mentioned in the protocols (Table S1), all default software parameters were kept.

### Child psychological and anthropometric outcomes

Supported by the novel tractography of multisystemic pathways, we subsequently explored a potential early-life neurological predispositions to psychological and metabolic disregulations and the likely WM bundles implicated. From the neonatal sample, 129 children were followed-up at 18 months (108/137 normal-weight vs 21/28 obesity exposed) and assessed on the Bayley Scales of Infant and Toddler Development III (Bayley, 2006) by dedicated study psychologists. Parents completed the Child Behavior Checklist (CBCL, Achenbach and Ruffle, 2000), the Early Childhood Behavior Questionnaire, very short form (ECBQ, Putnam et al., 2006) and the Quantitative Checklist for Autism in Toddlers (QCHAT, Allison et al., 2008). Outcomes captured here were the *Internalising* and *Externalising* scores from the CBCL, the temperamental traits for *Surgency*, *Negative Affect* and *Effortful control* from the ECBQ and the *receptive language*, *expressive language* and *cognitive* subscales of the Bayley’s. Child weight (kg) and height (cm) were also measured at the visit. WHO standardised z-scores for age and sex (weight-for-age and length/height for-age z-scores [WAZ and LHAZ]) were calculated with the R package *whoanthro*.

### Statistical Analysis

Regression path models were built with the FBA metrics (mean FD, mean FC, mean FDC) for each tract in each hemisphere as dependent variables. All FBA metrics were normalised for easier comparisons between tracts, and predictors were checked for normality. In Model 1 we studied cross-sectional development in the normal-weight-exposed control neonates only, across the 37-44 weeks PMA window and the effects of sex assigned at birth, neonatal birthweight centile (BWC at delivery, calculated adjusted for sex and age at birth by INTERGROWTH, Villar et al., 2014), total brain tissue volume (TBV cm^3^) and postnatal age in days (PN age), see Table S2 and Figure S5. In Model 2, the exposure variable (normal-weight/obese) and the interaction exposure*PMA was added and models run again. Models 1 and 2 accounted for covariate associations to improve quantification of main effects (sex, PMA, BWC, PN age and obesity exposure). In Model 3, we predicted 18-month psychological and anthropometric outcomes from the FD for each tract (adjusted for sex and PMA) and obesity exposure. Child outcomes were adjusted for sex and age at assessment and maternal years in education. All covariances between these outcomes were also included. We report fit indices with the following cut-offs for good model fit: the Root mean square error of approximation =<0.06 (RMSEA; Steiger, 1990) and the standardized root mean square residual (SRMR) <0.10 for good and <0.05 for very good fit (Steiger, 1990), comparative fit index (CFI) and Tucker-Lewis index (TLI) >0.95. All models were run in *Mplus v8.3* (Muthén and Muthén, 2017), using the robust maximum likelihood estimator (MLR) and the outputs integrated with the R package *MplusAutomation* (Hallquist and Wiley, 2018). Although exploratory, we also provide an FDR correction at p<0.05 in Model 3 analyses.

## Results

### Projections in the amygdala and temporal pole

Figure 1C and the accompanying video show the generated streamline bundles. The same protocols were used for homologous tracts on both sides of the brain and are provided in supplementary material Table S1. The neonatal ST, vAMP and Am-NAcc generated follow the topography reported in the adult brain, presented in Figure 2, and we were further able to present subcompartmental demarcation within the amygdala, where the tracts appear discreetly organised in the basolateral/central amygdala areas. The ST projections terminate preferentially inferior to those of the UF and vAMFP, as seen on the coronal slices (Figure 2A). The organisation of the ST, vAMFP, Am-NAcc and UF show little mixing. The UF brushes the amygdala lateral to the other two fibre bundles as it enervates the cortical areas of the temporal pole. The streamlines of the ST running caudal to the amygdala correspond to the centromedial and basolateral and lateral regions, mirroring the descriptions of this tract by Mori et al. (2017). On the axial and sagittal slices (Figure 2B and C), the topographical organisation anterior to posterior is as such: UF, Am-NAcc, vAMFP and ST.

**Figure 2:**
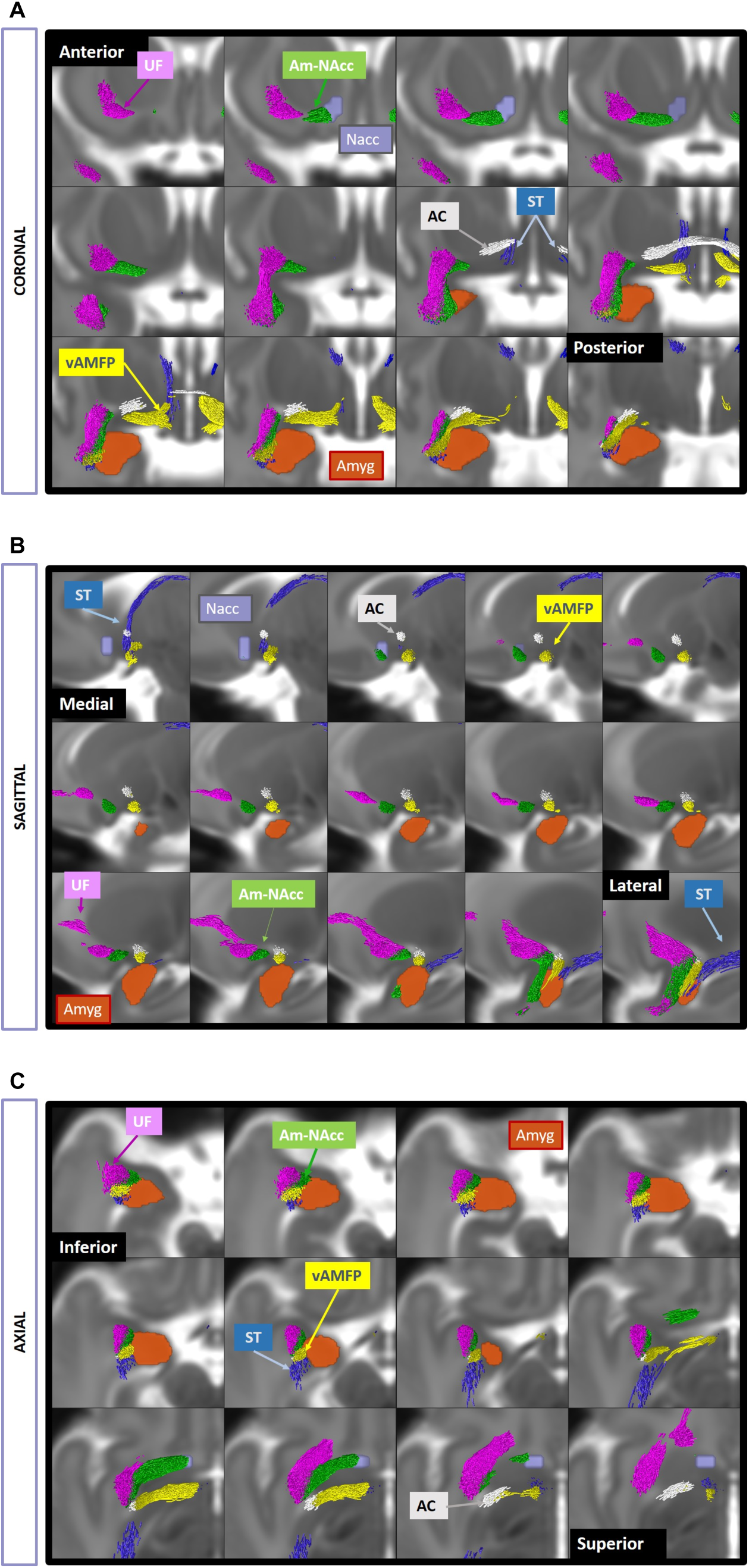
Right coronal (A), axial (B) and sagittal (C) sections through the temporal lobe and diencephalon demonstrate the topographical organisation of the Stria Terminalis (ST), Uncinate Fasciculus (UF), ventral amygdofugal pathway (vAMFP) and the Amygala-accumbens fasciculus (Am-NAcc).

### Projections through the hypothalamus

On their posterior approach, the DLF streamlines descending from the hypothalamus were specific to the medial flanks of the third ventricle (periventricular area), possibly the nucleus posterior hypothalami, superior to the bundles of the MFB at the level of the subthalamic nuclei and mammillary bodies. The MFB entered the lateral portion of the hypothalamus as described in the adult.

Of note in the left DLF were a few streamlines emminating further ventrally (Figure 3A), which we speculate are the cross-nuclei projections between the posterior nucleus into the ventromedial nucleus. We also observed some streamlines from the DFL joining into the MFB (Figure 3A, orange box).

**Figure 3:**
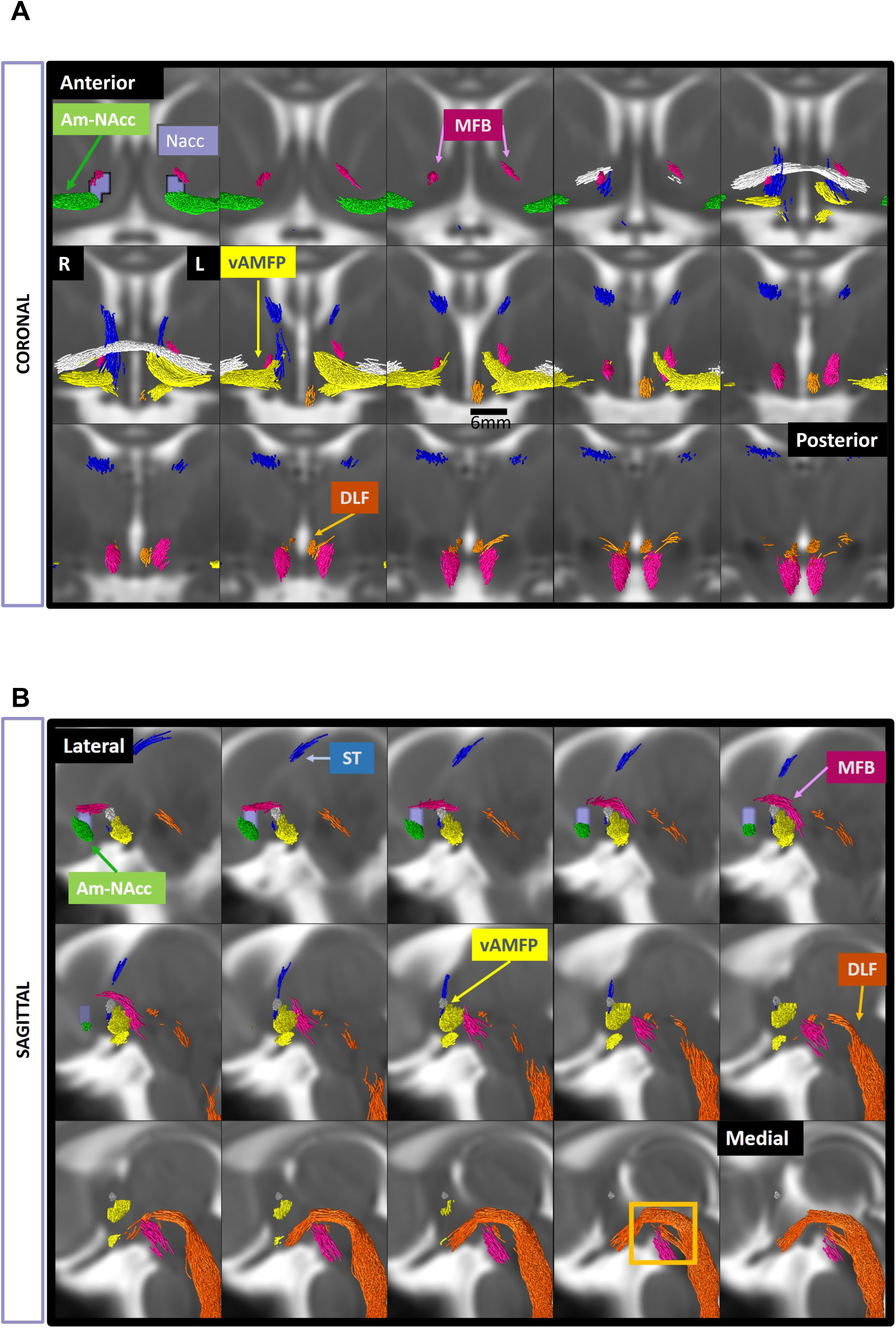
Coronal (A) and left sagittal (B) sections through the hypothalamus demonstrate the topographical organisation of the Stria Terminalis (ST), Dorsal longitudinal fasciculus (DLF), ventral amygdofugal pathway (vAMFP) and the Amygala-accumbens fasciculus (Am-NAcc) and Median forebrain bundle (MFB). In the sagittal plane, the yellow box may be the joining of the MFB and DLF fibres.

The vAMFP shows an approach posterior to the UF and Am-NAcc, inferior and parallel to the AC as it courses under the lentiform nuclei (internal globus pallidus), as appreciated on all the planes in Figure 2. Medially, the streamlines terminated at the anterior hypothalamic area. This follows the description of Kamali et al. (2016) in the adult but we find an additional split within the hypothalamus, i.e. of the fibres ascending towards the thalamic peduncle as described by Willis and Haines (2018). Unlike the description by Kamali et al. (2016), we made the distinction between the tract of the vAMFP and those fibres reaching the NAcc (Am-NAcc tract) which, on the axial plane, routed through the substantia innominata.

### Projections to the NAcc

The MFB joining the VTA to the NAcc showed bilateral differences in regards to the reported branches (superior and inferior to the Anterior Commissure). The right MFB was primarily coursing inferior whereas the left MFB streams were confined to the superior aspect i.e. through the inferior limit of the anterior limb of the internal capsule (Figure 3). In both MFBs the preferred termination corresponds to the shell of the NAcc. This is in line with histology (Haber et al., 2000).

The Am-Nacc streamline bundle leave the basolateral amygdala and showed a course through the Ansa lenticularis-pathway, which was described by Rusche et al. (2021) in their adult tractography. The termination points in the neonatal NAcc were primarily into the ventral NAcc, although Rusche et al. (2021) alluded to the central NAcc.

### Neonatal brain development in neonates born to mothers of normal-weight with an uncomplicated pregnancy

There were no differences in any maternal characteristics or delivery mode between males (n=77) and females (n=60). Girls were born later (40.09[1.01] vs 40.46[1.07], p=0.040) and, as widely reported, boys had larger birth head circumference (mean[SD]: 34.81 [1.43] vs 34.21[1.63] cm, p=0.022).

Figure 4 shows the correlation between WM tracts measures of micro (FD) and macrostructure (FC, FDC) across 37 to 44 week PMA. Figure 5 presents the standardized coefficients of the main effects from the regression path models (Figure S5B) which all showed good model fit (CFI >=0.99 ; TLI>=0.97; RMSEA >=0.06; SRMR >=0.05).

**Figure 4:**
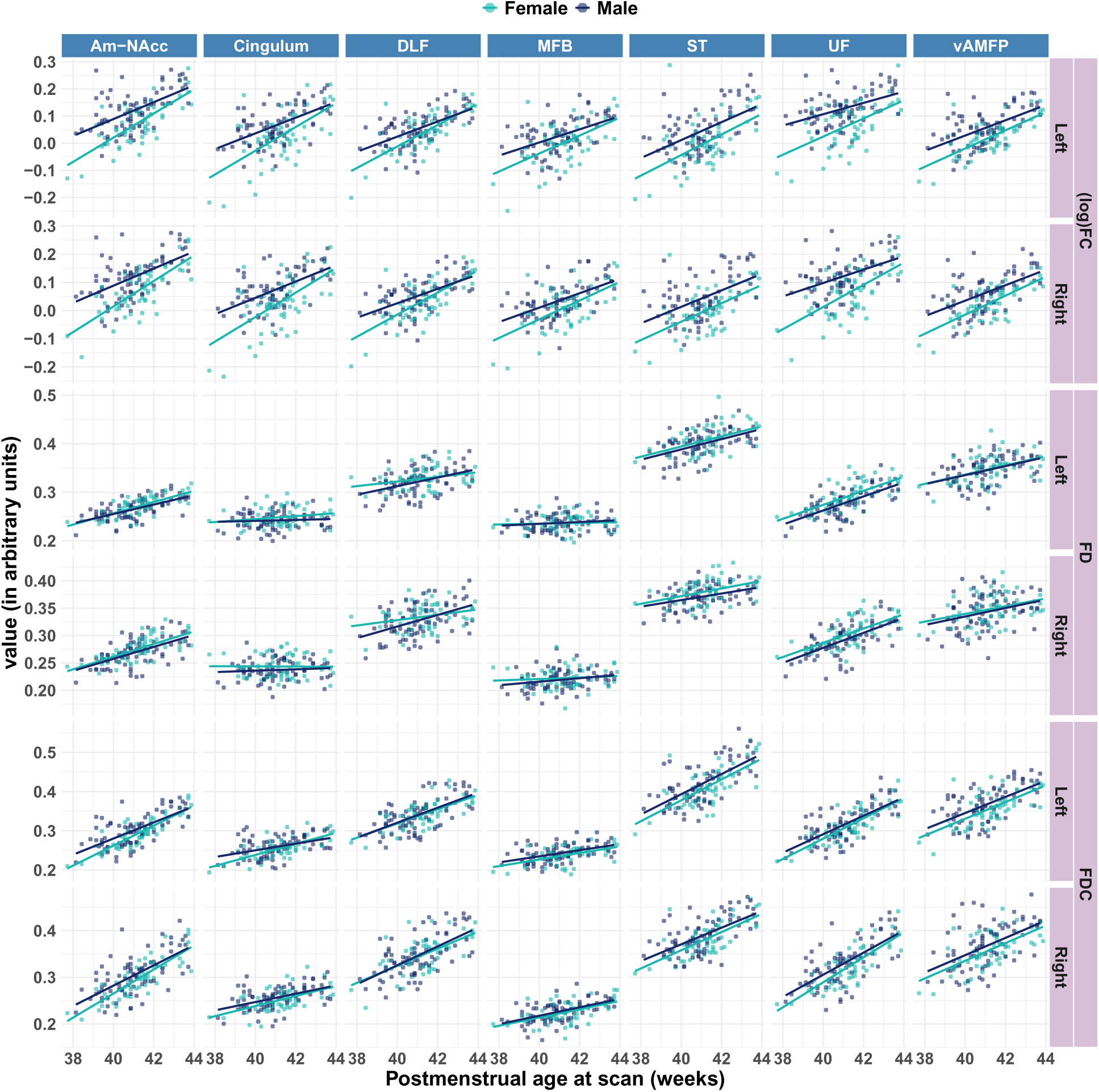
In normal-weight exposed neonates, correlations between postmenstrual age at scan and fibre density (FD), fibre cross-section (FC) and combined fibre density and cross-section (FDC). Am-NAcc: Amygdala-Nucleus Accumens tracts, DLF: Dorsal longitudinal fasciculus, MFB: Medial forebrain bundle, Stria Term.: Stria terminalis, UF: Uncinate fasciculus, vAMPF: Ventral amygdofugal pathway.

**Figure 5:**
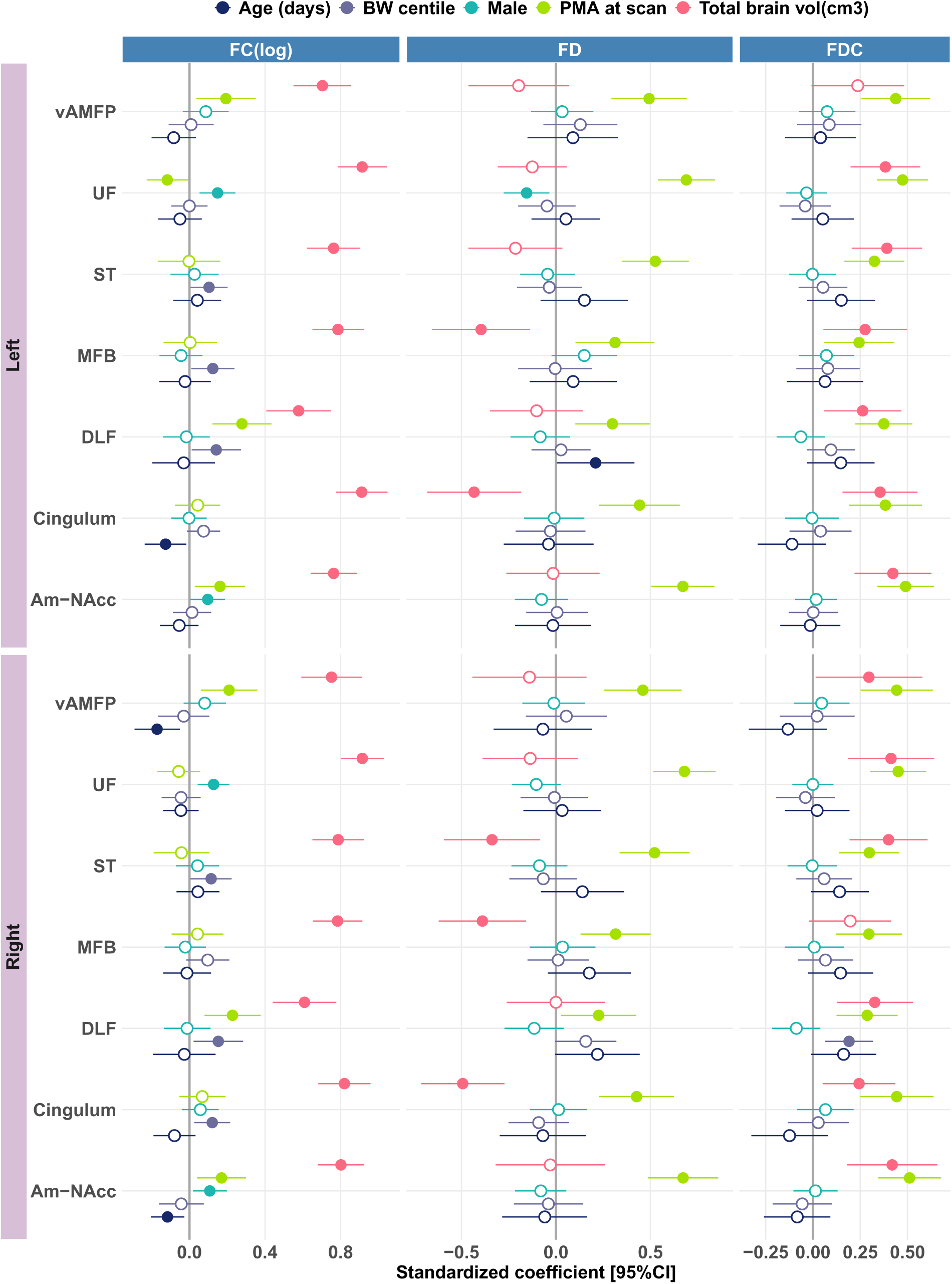
By path analysis, regression coefficients are presented for the effects of covariates on fibre density (FD), fibre cross-section (FC), fibre density x fibre cross-section (FDC) in each hemisphere, in 77 male and 60 female neonates of normal-weight women. Circles are filled if the 95% confidence interval excludes 0. Note the models included other regression paths as represented in the Figure S1C. DLF: Dorsal longitudinal fasciculus, MFB: Medial forebrain bundle, ST: Stria terminalis, UF: Unicinate fasciculus, vAMPF: Ventral amygdalofugal pathway.

Holding other covariates constant, FC grew with PMA in the vAMP, DLF and Am-NAcc bilaterally, but constricted with PMA in the left UF and did not change in the bilateral cingulum, MFB and ST. Males had larger FC in the bilateral UF [standardized beta [95%CI] Left: 0.15[0.054 to 0.24], p=0.002, Right: 0.13[0.044 to 0.21], p=0.003) and bilateral AM-NAcc (Left: 0.097[0.006 to 0.19], p=0.036, Right:0.11[0.019 to 0.20], p=0.018). TBV expectedly had the largest positive effect on FC of all the covariates across all the tracts, highest in the cingulum. Infants of higher BWC had larger FC in bilateral ST, bilateral DLF, left MFB and right cingulum. Postnatal days outside the womb had a negative effect (i.e. constriction) on the left cingulum, right AM-NAcc and right vAMFP.

FD increased with increasing PMA in all tracts, with the largest effects seen in the bilateral UF and Am-NAcc (Figure 5). Males had lower FD in the left UF (–0.15[-0.28 to –0.035], p=0.012). TBV was negatively associated with the FD in the bilateral cingulum, bilateral MFB and right ST. Postnatal age in days was positively associated with the FD of the DLF. BWC had no effect on FD.

PMA at scan and TBV associated with FDC and infants of larger BWC had larger FDC in the DLF.

### Comparisons by obesity exposure

Obesity and obesity*PMA were then added as predictors in the path models. The scatterplots of the correlations between FBA metrics and PMA with exposure as the grouping variables are shown in Figure S6. Path analyses on FBA metrics showed model fit were good (CFI> 0.96, TLI: 0.93 to 0.97, RMSEA:0.062, SRMR: 0.056 to 0.060). There was no main effect of the exposure (normal-weight vs obesity) on any of the tracts. There was a significant negative interaction of exposure*PMA on the FD in the bilateral Am-NAcc tract (Right: –0.57[-0.92 to –0.21, p=0.002]; Left: –0.57[-1.10 to –0.03], p=0.038) and the right UF (–0.47[-0.84 to –0.10],p=0.013), see Figure S7.

### Associations with outcomes at 18 months

We explored whether neonatal fibre density and obesity exposure predicted psychological and anthropometric outcomes in the children at 18 months. Figure 6 represents the beta coefficients. Results show that obesity-exposed children had higher WHAZ (0.23[0.05;0.41],p=0.012) and LHAZ (0.2[0.02;0.38], p=0.020) and lower cognitive scores (–0.29[-0.45;-0.13], p<0.001) than non-exposed at 18 months. In the whole sample, neonatal FD of the bilateral Am-Nacc positively predicted Surgency traits (left= 0.22[0.07;0.36],p=0.004; right= 0.19[0.03;0.34], p=0.022), and higher right FD of Am-NAcc predicted lower effortful control (–0.17[-0.34;0.00],p=0.046). The higher FD in bilateral cingulum associated with higher QCHAT score (right=0.22[0.06;0.39],p=0.008; left=0.28[0.12;0.44], p<0.001). Higher FD in the left cingulum predicted higher Negative Affect score (0.21[0.06;0.37], p=0.008) whereas higher FD in the right cingulum predicted lower Receptive Language score (–0.18[-0.36;0.00], p=0.044). Higher FD in the bilateral DLF predicted higher Expressive Language score (right=0.23[0.08;0.38], p=0.002; left=(0.18[0.03;0.33], p=0.023) and the right DLF predicted higher Receptive Language (0.28[0.13;0.43], p<0.001) and Cognitive score (0.19[0.02;0.36],p=0.028) and lower QCHAT score (–0.19[-0.38;-0.01], p=0.044). For the MFB, only the right positively predicted Surgency (0.19[0.03;0.36], p=0.021). FD of the right vAMFP associated with higher Negative Affect (0.16[0.01;0.32], p=0.038). The neonatal FD of the ST was not predictive of any 18-month outcomes. None of the neonatal WM tracts FD predicted 18-month anthropometry. However, child WAZ correlated significantly with higher Internalising (0.22[0.09;0.36], p=0.006), Externalising (0.28[0.12;0.45], p<0.001) and Surgency traits (0.20[0.02;0.38],p=0.030) and with lower Effortful Control (–0.22[-0.38;-0.06],p=0.006). LHAZ correlated with Surgency only (0.23[0.07;0.4], p=0.005). Table S3 provides parameters results, p-values and fdr-corrected p-values for the paths tested which show effects of the Cingulum on QCHAT and DLF on Receptive Language and Cognition survived correction for multiple comparisons.

**Figure 6:**
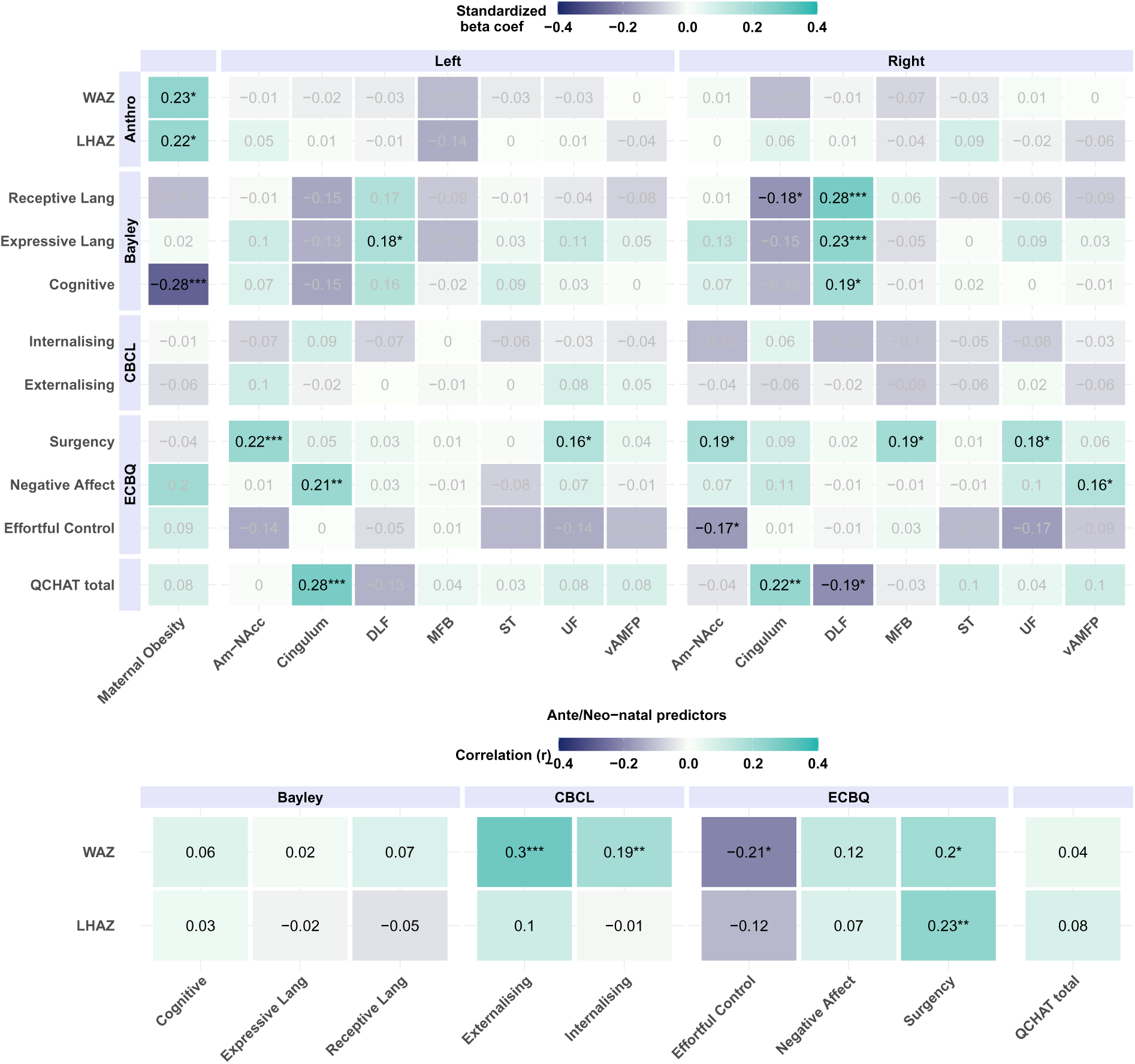
Results from path regression models predicting child psychological and anthropometric outcomes at 18 months, adjusted for child age at assessment and sex and maternal education. A: Beta coefficients of bilateral fibre density and maternal obesity group as predictors. B: Correlations between 18-months outcomes included in the paths models. Stars denote significance levels *p<0.05,**p<0.01 and *** p<0.001. CBCL: Child Behavioural Checklist, DLF: Dorsal longitudinal fasciculus, ECBQ: Early Childhood Behavior Questionnaire, LHAZ: sex-adjusted length or height-for age z-score, MFB:Medial forebrain bundle, QCHAT: Quantitative Checklist for Autism in Toddlers, ST:Stria terminalis, UF:Uncinate fasciculus, VAMPF:Ventral amygdalofugal pathway, WAZ: sex-adjusted weight-for-age z-score.

## Discussion

This study contributes to the field of neuroimaging and epidemiology in several ways. First, the neonatal Amygdala-Nucleus Accumbens fasciculus (Am-NAcc), dorsal longitudinal fasciculus (DLF), medial forebrain bundle (MFB), stria terminalis (ST) and ventral amygdalofugal pathway (vAMFP) were delinieated for the first time using tractography. Second, we demonstrated the topographical organisation of these pathways within the hypothalamic and amygdala subregions, which has not been characterised previously *in vivo* nor at this age. Third, FBA is a method sensitive to voxel-wise tissue microstructure of density and cross-section of *distinct* bundles and through path analysis, we found heterogeneity in WM growth rate across 6 weeks postpartum and quantified the effects of TBV, BWC and sex. Fourth, we compared WM structure between offspring of normal-weight and obese women. This suggested a negative interaction between maternal obesity and PMA for FD in the bilateral Am-NAcc bundle and the right UF. Lastly, we identified potential neurological pathways which, independently or in combination, may explain the relationship with child temperamental outcomes and their relationship with body size as early as 18 months.

In terms of tractography outputs, given the smaller brain size which could prevent such spatial sensitivity, we were surprised to find the topology of most tracts within each ROI concurred with previously reported histology in the adult. For example, the MFB streamlines crossed the lateral hypothalamus and terminated in the NAcc distinctly from those of the Am-NAcc streamlines in respect to the NAcc core and shell. Additionally, we showed the distinction between the bundles of the Am-NAcc and those of the vAMFP within the hypothalamus, although they share the temporal lobe portion emanating from the amygdala. Otherwise, organisation of the vAMFP and Am-NAcc in relation to the UF shown here in the neonate agrees with previous adult reports (Folloni et al., 2019). Rusche et al. (2021) provided Nacc derived probabilistic tractography and ours agrees with their delineation of the VTA-NAcc path (i.e. MFB) and Am-NAcc path. However, our study further demarcates the spatial organisation of these tracts in relation to each other within the neonatal amygdala and the hypothalamus. Nevertheless, we obtained a right vs left MFB lateralisation (superior vs inferior branch) which we consider to be a false negative effect since these two branches have been generated previously by tractography in the adult (Bracht et al., 2015; MacNiven et al., 2020). As for the ST, it is visually thin according to adult histology and may be still going through fasciculation in our sample. Importantly, the DLF carries the autonomic transmission between the vagus nerve to/from the hypothalamus. When the whole hypothalamus served as the included region of interest in the protocol, we showed primary bilateral terminations of the DLF were in the (sympathetic) posterior hypothalamic area, but also with a smaller portion/branch terminating towards the ventromedial area, corroborating research in older human and animal studies(Goel et al., 2024).

The data presented here shows that WM density increases in the perinatal period but not at the same rate across all bundles. We observed that the DLF and MBF had the lowest rate of FD change. This may reflect the medial-to-lateral direction of neuronal expansion in early development and earlier maturity being reached in these tracts. Of note, as a voxel-wise measure of axonal matter, FD cannot distinguish between a change in axonal count or axonal diameter and is not directly sensitive to myelin. FD can also be influenced by the other components in the extra-axonal space surrounding a fibre bundle(Dhollander et al., 2021). However, in regards to the DLF, autonomous neonatal fibre count may be fixed at birth (Sachis et al., 1982).

Differences in brain anatomy between the sexes such as in this study have been reported (Kim et al., 2017) although not consistently in the term born neonatal WM (Akazawa et al., 2016; Dean et al., 2017). Comparisons to previous studies are difficult since others have included large WM tracts or regions and different metrics of assessing WM development than FBA. Nevertheless, others have found growth rate of some WM regions show sex dimorphism in later childhood development (Deoni et al., 2012) and larger temporal cortex (medial) in the male neonate (Knickmeyer et al., 2014) which appears to be conserved throughout the adult lifespan (Lotze et al., 2019). Interestingly, amygdala volume growth trajectories are highly dimorphic between 5 and 25 years of age which is particularly prominent in the centromedial nuclear group of the amygdala (Fish et al., 2020). Therefore, the larger UF and Am-NAcc tract fibre cross-sections in male neonates showed in this present study could be in agreement with these previous findings since the two tracts serve the temporal/amygdala connectivity to frontal areas. This aligns with findings on the frontal and temporal regional maturation in an overlapping sample using morphometric similarity networks (Fenchel et al., 2020).

From the group of 28 neonates exposed to obesity *in utero*, our results suggest that, in both sexes combined, those neonates followed a slower FD growth trajectory compared to infants of normal-weight women in the bilateral Am-NAcc and the right UF. Comparisons with recent literature are limited since offspring MRI are acquired differently, using tensor-based models in groups scanned in childhood or adulthood and so potentially influenced by residual postnatal exposures (Parsaei et al., 2024). From those, Ou et al. (2015) detected wide-spread differences in fractional anisotropy in a small sample of obesity-exposed neonates. In preterm-born neonates with no radiological abnormalities, Lee et al. (2023) found lower axial diffusivity in the UF of those exposed to high maternal BMI and Rasmussen et al. (2023) showed that neonatal mean diffusivity is lower in the hypothalamus as a whole following obesity exposure. These studies did not present outcome measures to infer on longer term impact.

### Early origins of psychobehavioural and anthropometric outcomes

In this present study we found that antenatal obesity exposure predicted higher body morphology at 18 month and lower cognitive outcomes. Although, we could not find a direct effect of neonatal WM density on anthropometry we did find that neonatal FD in the two same tracts vulnerable to obesity exposure (i.e. UF and Am-NAcc) predicted higher Surgency traits and lower Effortful control. Furthermore, we also found that the right superior MFB branch predicted Surgency at 18 month which partly agrees with previous work which demonstrated lower FA in the inferior MFB branch was associated with higher impulsivity (MacNiven et al., 2020) and the superior branch mean FA negatively associated with hedonistic capacity (Bracht et al., 2015). Importantly, we found that higher Surgency and lower Effortful Control (and higher Internalising/Externalising scores) were correlated with higher weight-for-age z-scores. Previously, Surgency has been described as a approach/reward-seeking temperament including high appetitive drive and obesogenic eating behaviour without hunger in preschoolers (Leung et al., 2014). Therefore, addressing the primary aim of this work, we found support for the hypothesis of an overlapping neurological underpinning between psychological and anthropometric phenotypes i.e. a risk to higher weight-for-age is mediated by a reward-seeking temperamental profile of early neurological origin which drives a potential for larger anthropometry and that deviation in development of these WM pathways may occur through gestational exposure to maternal obesity. Such predisposition could be reflected in children with higher emotionally-driven reward needs seeking higher food intake (Am-NAcc, UF) and/or present dysregulation in energy metabolism directly and indirectly via the action of the hypothalamus (MFB, vAMFP). Antenatal biological mechanisms implicated in obesity exposure could include an alteration of both the fetal inflammatory and neurodevelopmental transcriptome profiles, directly or indirectly via placental dysfunction, and more specifically disruption of the patterning genes involved in dopamine neuron differentiation (Ma et al., 2014; Money et al., 2018).

Our lack of association of the neonatal DLF FD and toddler anthropometry in this study may represent a distinct functional response path of the DLF into its receptive hypothalamic subregion of the ventromedial hypothalamus. Alternatively, the FD metric may not be reflective of this DLF function. It is possible that the DLF implication in anthropomorphy through orexigenic/anorexigenic or microbiotic signals is shaped, not by axonal density at birth, but rather the myelination process, chemical substrates and neurotransmitter release within the gut-brain-axis. These processes could be either disrupted by *in utero* exposure or by external factors later in postnatal development (gut pathogens invasion, diet quality). In contrast, the MFB would act upon the dopaminergic pathway through the lateral hypothalamic nuclei to predispose to a reward-seeking temperament.

However, based on the above observations of the associations between neonatal WM tracts and outcomes, we speculate further to have also shown partial support for the structural underpinning linked to the Polyvagal Theory (Mulkey and Plessis, 2019). The theory suggests that the autonomic nervous system directs and modulate the extent of emotional, social and behaviour responses via neurophysiological substrates (Porges, 2007a). It proposes, for example, that expressive communication and emotionally-elicited behaviours hinges on vagal inhibition or activation and its interactions with other regulatory system such as the HPA-axis (Porges, 2007b). Additionally, it has been shown that sensory over– and underresponsiveness are linked to autism (Jung et al., 2021). In this study, the DLF does not appear to be implicated in the temperamental or internalising/externalising outcomes but rather, higher fibre density explained higher overall cognitive and language ability and also lower autistic traits (QCHAT scores). In the cingulum, in contrast, denser fibre bundles associated with higher QCHAT and lower receptive language and higher negative affect scores. The DLF, transmits autonomous signals between the hypothalamus and the vagal preganglionic fibres and thus directly supports the arousal reactivity of an individual. The cingulum is connected widely to sensorimotor regions and is necessary for sensorimotor coordination, arousal and emotional dysregulation (Caruana et al., 2018; Hung et al., 2020). We speculate therefore that higher fibre density of the DLF reflects a more mature, and perhaps “efficient”, vagal capacity which may offer the scaffolding needed to engage and attend in cognitive and language processing and regulate emotional reacitvity. This autonomic system may then be counterbalanced by the limbic-reward response (via cingulum, MFB, UF, Am-NAcc, vAMFP shown here to associate with Surgency and/or Effortful Control), where denser WM may reflect a vulnerability at birth, by genetic liability or suboptimal *in-utero* exposure (i.e. in our case the influence of obesity on Am-NAcc and UF development), and predicts more adverse affective/sensory outcomes in the longer term.

Despite methodological strengths, our work presents some limitations. As mentioned, comparing tractography results and quantifications with those used in the adult population is not straightforward and postmortem histology in the neonate is scarce for a ground truth evaluation. Importantly, the brain in the perinatal period is dynamically changing and accounting for processes such as inter-subject fasciculation, axonal growth and myelination is challenging. We prioritised the “core” body of each fibre bundle where the macro/microstructure of the axonal bundle is assumed to be the least sensitive to this variability (e.g. temporal stem for the uncinate fasciculus) compared to the distal cortical terminations. This should quantify by proxy the direct transmission capacity between two cortical/subcortical areas, represented by the FC, FD, FDC within this core. Calculating their mean over a core tract aimed to reduce issues of partial volume effect within voxels at the periphery of the tract but may misrepresent differences along the tracts. Finally, we make some assumptions, e.g., that our DLF tractography is specific to the vagal transmission, which we cannot confirm.

Beside an imbalance in ethnic representation between the exposure groups, we acknowledge that we only had a convenience sample to explore the impact of antenatal obesity exposure and our analyses may have been underpowered to detect further effects at 18 months. Moreover, here we excluded obstetric complications, preterm births and brain abnormalities which may remove the mediating or additive risks on the causal path to adverse offspring long term health outcomes. Nonetheless, our findings emphasize that obesity alone in pregnancy was still associated with structural brain difference once these mediators and additional comorbidities are removed. Additionally, other effects of obesity exposure on brain microstructure may be sex-dependent or differences in WM between these groups appear later in development (Akazawa et al., 2016). Finally, a potential role for genetic predispositions and maternal mental health in these obesity-exposure effects should not be ignored (Wilson et al., 2018), nor their interaction with maternal antenatal diet, infection and inflammation (Sigurdardottir et al., 2022) or, postnatally, parental use of food to soothe child temperament (Stifter and Moding, 2018).

In summary, state-of-the-art dMRI acquisition and processing can deliver topographically sensitive delineation of WM tracts in the neonatal brain, with a level of detail and specificity only previously elucidated from animal retrograde/anterograde histology and postmortem examinations. Longer term exploration of the functional relevance of the sex differences and effect of obesity exposure observed in WM development and child outcomes is warranted. The potential utilisation of the novel protocols devised in this study is substantial, especially in defining potential early brain biomarkers among infants at higher likelihood of neurodevelopmental conditions such as those born preterm or with familial history.

## Data availability

The datasets used in the current study are available in the dHCP repository http://www.developingconnectome. org/data-release/second-data-release/

## Supporting information

GIF of Tracts

## Acknowledgement

We acknowledge and are thankful for all the parents of the neonates who contributed to this study and the staff at St Thomas Hospital. The dHCP was funded by the European Research Council under the European Union Seventh Framework Programme (FP/2007-2013)/ERC Grant Agreement No. 319456. JNS was funded by a PhD studentship from the National Institute for Health Research (NIHR) Biomedical Research Centre based at Guy’s and St Thomas’ NHS Foundation Trust and King’s College London. The authors received funding from the Innovative Medicines Initiative 2 Joint Undertaking under grant agreement No 777394 for the project AIMS-2-TRIALS. This Joint Undertaking receives support from the European Union’s Horizon 2020 research and innovation programme and EFPIA and SFARI, Autistica, Autism Speaks. The views expressed are those of the authors and not necessarily those of IHI-JU2. The funders had no role in the conceptualisation of this study, nor the development of this publication. This paper also represents independent research part funded by the infrastructure of the National Institute for Health Research (NIHR) Mental Health Biomedical Research Centre (BRC) at South London and Maudsley NHS Foundation Trust and King’s College London. The views expressed are those of the author(s) and not necessarily those of the NIHR or the Department of Health and Social Care.

## Authors contributions

JNS processed and analysed the data, interpreted results, created the figures, wrote the first manuscript.JDT processed the data and interpreted results. DB, AB, DC, JH, LCG processed the data. AC, CN supported and collected 18-month outcome data. JH provided funding. GM, LP and MR supervised, interpreted the results. All authors reviewed and approved the manuscript.

