## Supplementary figures and images for "*In vivo* tractography of human neonatal white-matter pathways underlying hypothalamic and reward functions to study predispositions to neurodevelopmental conditions and obesity"

### GIF of Tracts

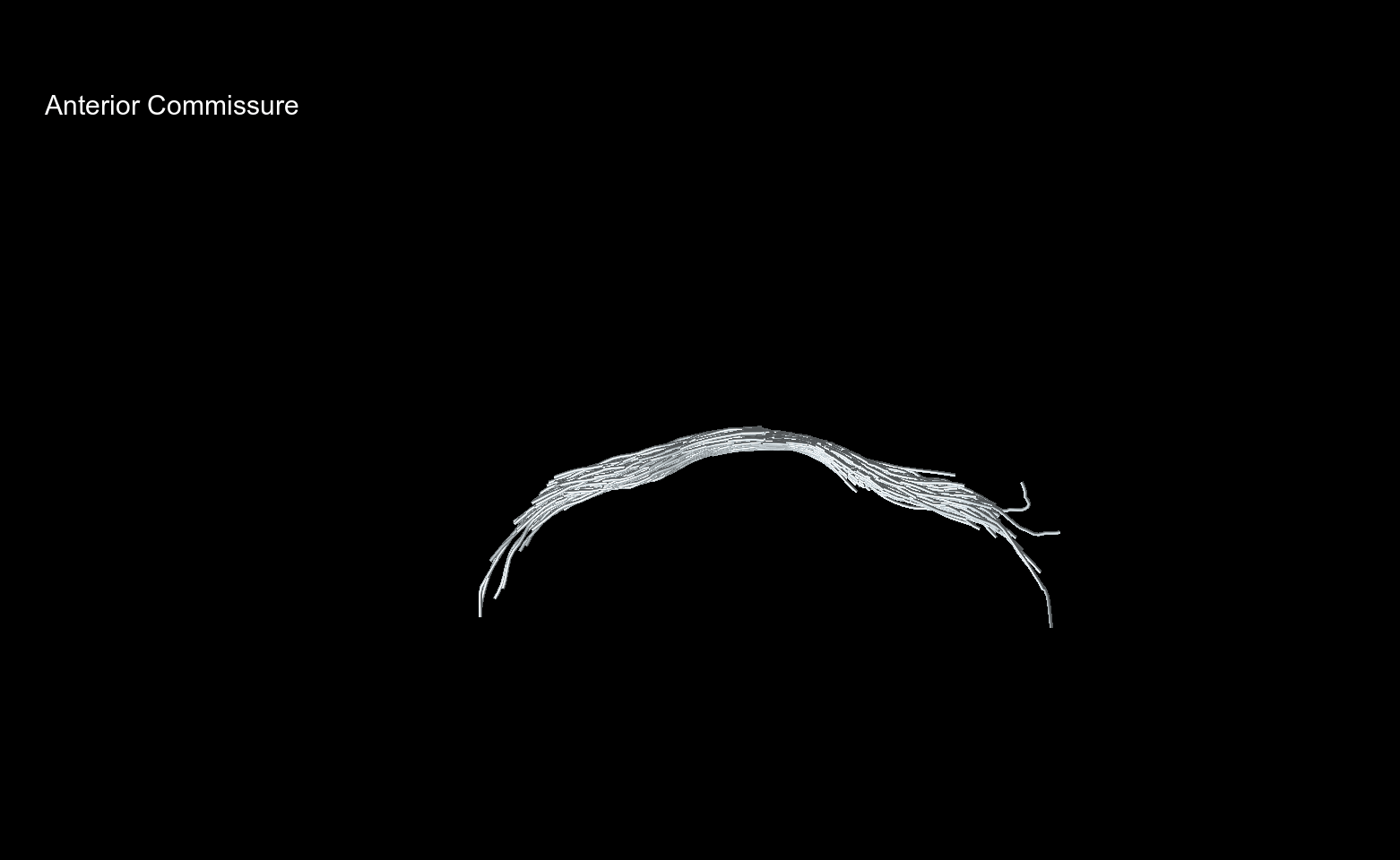
