## Supplementary File for "*In vivo* tractography of human neonatal white-matter pathways underlying hypothalamic and reward functions to study predispositions to neurodevelopmental conditions and obesity"

### Supplementary: '*In vivo* tractography of the neonatal white-matter pathways underlying hypothalamic and reward function to study early predisposition to neurodevelopmental conditions and obesity.'

Sigurdardottir et al.

#### Tracts of interest

Note that, although coursing through the vAMFP, the amygdala to NAcc tract was differentiated and referred to as Am-NAcc (also known as Amygdalo-accumbens fasciculus), whereas the amygdala to hypothalamus tract is referred to as the vAMFP.

**Amygdala-NAcc pathway (Am-NAcc)**

The Am-NAcc is also known as the amygdaloaccumbens fasciculus (1). It courses through the temporal section of the vAMFP from the anterolateral amygdala, but separates from these fibres reaching the hypothalamus, to target the NAcc anteriorly, preferentially to its "shell", the ventromedial portion of the NAcc. This path is otherwise immediately adjacent to the UF.

**Cingulum**
The cingulum is an association tract which is contained within the cingulate gyrus dorsally and the parahippocampal gyrus as it curves around the corpus callosum into the temporal pole. In this study the delineation was focused on the long curving fibres and not the short radiation fibres joining the cortex. It runs ventral to the subiculum of the hippocampus and does not contact the fornix, which must be differentiated accurately in the *in vivo* tractography.

**The Dorsal longitudinal fasciculus (DLF)**

The DLF originates from the medial and periventricular hypothalamic zones, passes through the periaqueductal grey matter (PAG) and caudal medulla oblongata and which transmit to neurons of the vagus nerve at (cranial nerve X).

**Medial Forebrain Bundle (MFB)**

The MFB associates the VTA to the NAcc through the lateral hypothalamus, the infero-medial branch of the MFB. Identified in the human is the superolateral branch from the VTA which accesses the NAcc through the inferior portion of the anterior limb of the internal capsule and superior of the AC, briefly mingling with the anterior thalamic radiation (2–5). Some descending MFB fibres mix with the DLF in the brain stem but only the segment VTA-NAcc was segmented here.

**Stria Terminalis (ST)**
The ST originates from the corticomedial amygdala (6) and terminates in the bed nucleus of ST (BNST), the hypothalamus (anterior and medial preoptic area) and septal nuclei. These fibres loop around the medial caudate nucleus, associating with it within the grove with the thalamus (i.e. caudothalamic groove) and laterally parallel with the fornix. The BNST lies dorsal to the AC, lateral to the columns of the fornix and medial to the thalamostriate vein (7). Corticomedial fibers of the amygdala pass through this pathway to reach the NAcc (8,9)

**Uncinate Fasciculus (UF)**
This association tract takes the shape of a hook from the anterior temporal pole, through the temporal stem, entering the extreme and external capsule and terminates in the orbitofrontal lobe.

**Ventral Amygdalofugal Pathway (vAMFP)**

Fibres in the basolateral nuclear group and central nuclei of the amygdala run medially through the substance innominata and substantia perforata anterior, fanning out into the thalamic peduncle and joining the diagonal band of Broca. In the hypothalamus it reaches the lateral preoptic area (9) and more anteriorly the septal nuclei (10) and the NAcc (11,12).

### 1 Exclusion criteria

From 962 participant in the dHCP database at the time of enquiry, we excluded non-singleton pregnancies (110), participants with missing obstetric outcomes (208) and missing BMI (39) and the following antenatal diagnoses : Obstetric cholestasis (6) ,preeclampsia (21), **H**amolysis, **E**levated **L**iver enzymes and **L**ow Platelets (HELLP) (1), pregnancy induced hypertension (25) and gestational diabetes mellitus (7). From the 545 remaining, 142 had no dMRI data available. 111 of 403 born were born outside the range 37-42 weeks gestational age and 89 women were BMI <18.5 or 25 to <30 kg/m$^2$thus excluded. 174 and 29 pregnancies of normal-weight (18.5 to 25 kg/m$^2$) and obese women (>=30 kg/m$^2$ ), respectively, remained. We excluded infants with apgar score below 7 at 5min (or missing) and who were admitted to the NICU. We reviewed the dMRI quality blinded to groups and the presence of anatomical imaging (T1w or T2w) which excluded 11 more for poor quality.

The radiological reports were reviewed blinded to maternal BMI and participants were excluded if any of the following was noted: germinal matrix hemorrhage (1), intraventricular hemorrhage (1) and bilateral temporal horn cyst (1). Many infants had WM punctate lesions (WMPL), so each scan was reviewed individually and the number of lesions and their location was noted. We chose to exclude scans with more than 4 lesions on the assumption of a potential wide-spread effect on our analysis and inference. In the remaining infants with WMPL, none were within our regions or tracts of interests. Infants with significant congenital abnormalities (e.g. cardiac defect) were excluded (3).

### 2 Template generation and fixel masks

In order to generate tracts of interest, a FOD template was produced from 20 randomly selected obesity-exposed and 20 controls (*population_template*). Next, all the other subjects FOD maps in native space were registered (*mrregister*) to the template FOD space which produced subjects-to-template and template-to-subject warps. The subject FOD masks in native space were then warped to FOD template space using these warps and then their intersection (*mrmath -min*) was used to create the FOD template mask.

Subsequently a fixel mask was generated (*fod2fixel*) using the FOD template and the FOD mask with a threshold of 0.06 on the peak amplidtude of positive FOD lobes, from which quantitative analyses of fibre density and fibre crossing could be performed. Next, all subject FOD maps were warped to FOD template without reorientation applied (*mrtransform*). Thereafter, a whole brain fixel mask was generated for each subject FOD in FOD template space (*fod2fixel*), whereby the number and orientation of fixels per voxel was obtained but also the corresponding per-fixel apparent fibre density (AFD, now *fd*). Next, a re-orientation of fixels of all subjects in template space relying on the warps was performed (*fixelreorient*). After, with all subject FODs spatially aligned to the FOD template, the *fixelcorrespondence* command allowed for matching every fixel across subjects. Lastly the fibre cross-section (fc) metric was computed to estimate an index of morphological differences between subjects using the warps employed in the earlier registration(*warp2metric*). This fc was log-transformed as per recommendations. Finally, the density and cross-section metric (fdc) was computed as the product of fd and fc.

Following tractography (see below) fixel masks were generated from each tract generated in the population template (*tck2fixel*). After visual inspection of the masks, streamline-per-voxel thresholds were applied to maintain only the core of each bundle (see Figure S3) and the mean FD,logFC and FDC were extracted for each subject for each tract.

### 3 Anatomical template

The subject T2 anatomical images from the dHCP dataset are reconstructed to a 0.5mm isotropic resolution. In order to produce a T2w anatomical template aligned in the FOD template space we obtained the T2w images in native space (co-registered with diffusion data) and these T2 anatomical images were warped to FOD template space using the warps generated in the subject FOD-to-template FOD registration and using the FOD template mask previously upsampled to 0.5mm resolution as the registration template.

The subject ROIs derived from the atlas was in subject space at 0.5mm isotropic resolution and the FOD template at 1.5mm so some visual checks and manual adjustment were necessary to avoid partial-voluming. Anatomical structures manually drawn were the hypothalamus (with mamillary bodies), NAcc and VTA. Those derived from the anatomical automatic labelling (AAL) atlas adapted to the neonatal dHCP data (13) were the amygdala, hippocampus, the anterior temporal lobe (medial and lateral) and the mid and orbitofrontal WM.

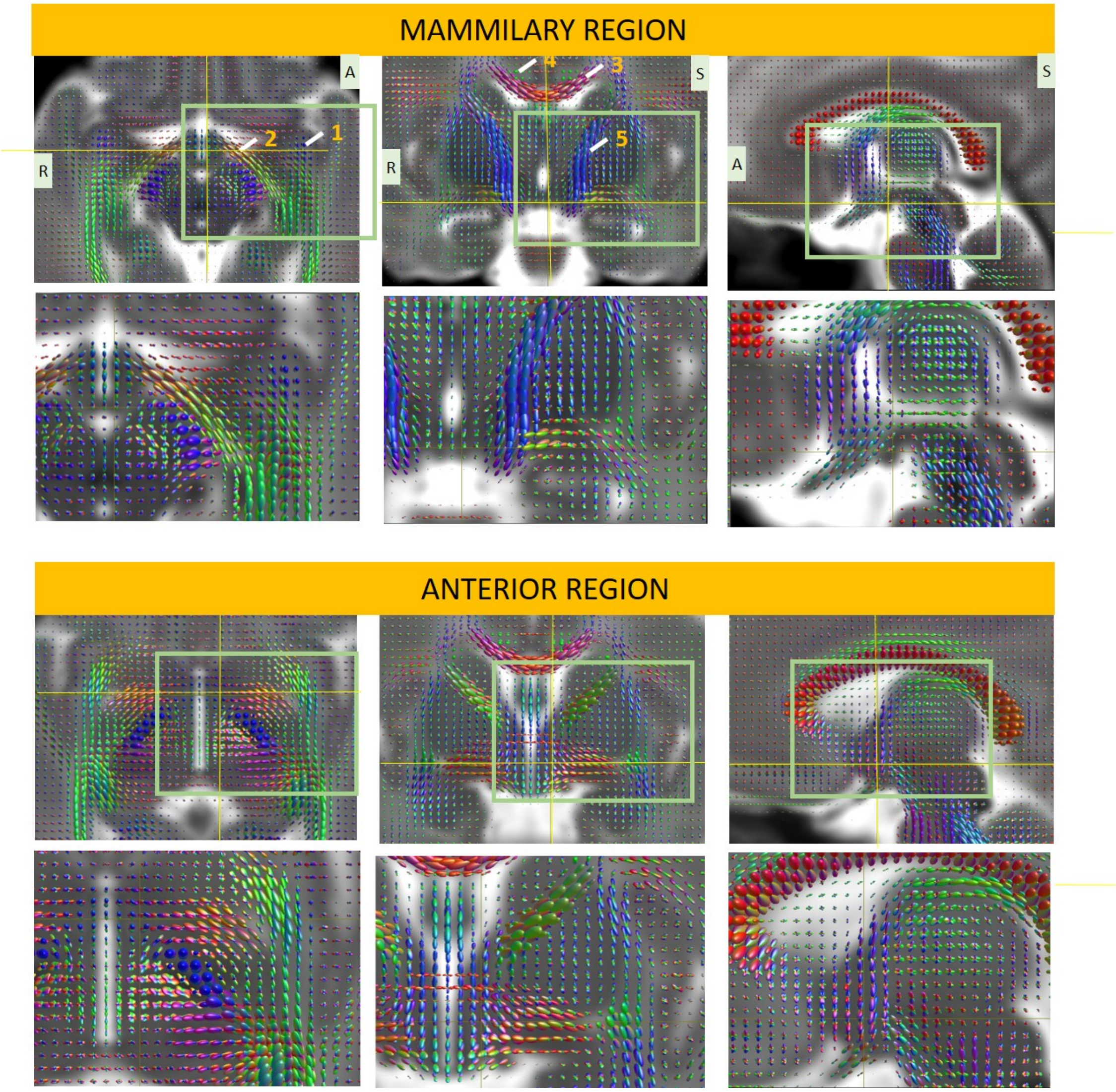


Figure S1: The population FOD template is overlaid on the T2 template at the anterior (anterior commissural) region and mammilary region of the hypothalamus. Bottom rows are zoomed in boxes from the top rows. The maps are presented in axial (first column), coronal (middle) and sagittal (right) views. Yellow lines are cross-hairs provided for orientation. FOD lobes are sized by peak amplitude and directions are color coded, red: left-right, blue:superior-inferior and green: anterior-posterior orientations. Note how some groups of s underly WM bundles easily distinguishable in the neonatal brain i.e. 1: ascending temporal stem of the uncinate fasciculus, 2:optic tract, 3: corpus callosum, 4:cingulate, 5: corticospinal tract. Note the T2 template is in 0.5mm isotropic resolution and the FOD template map at 1.5mm.

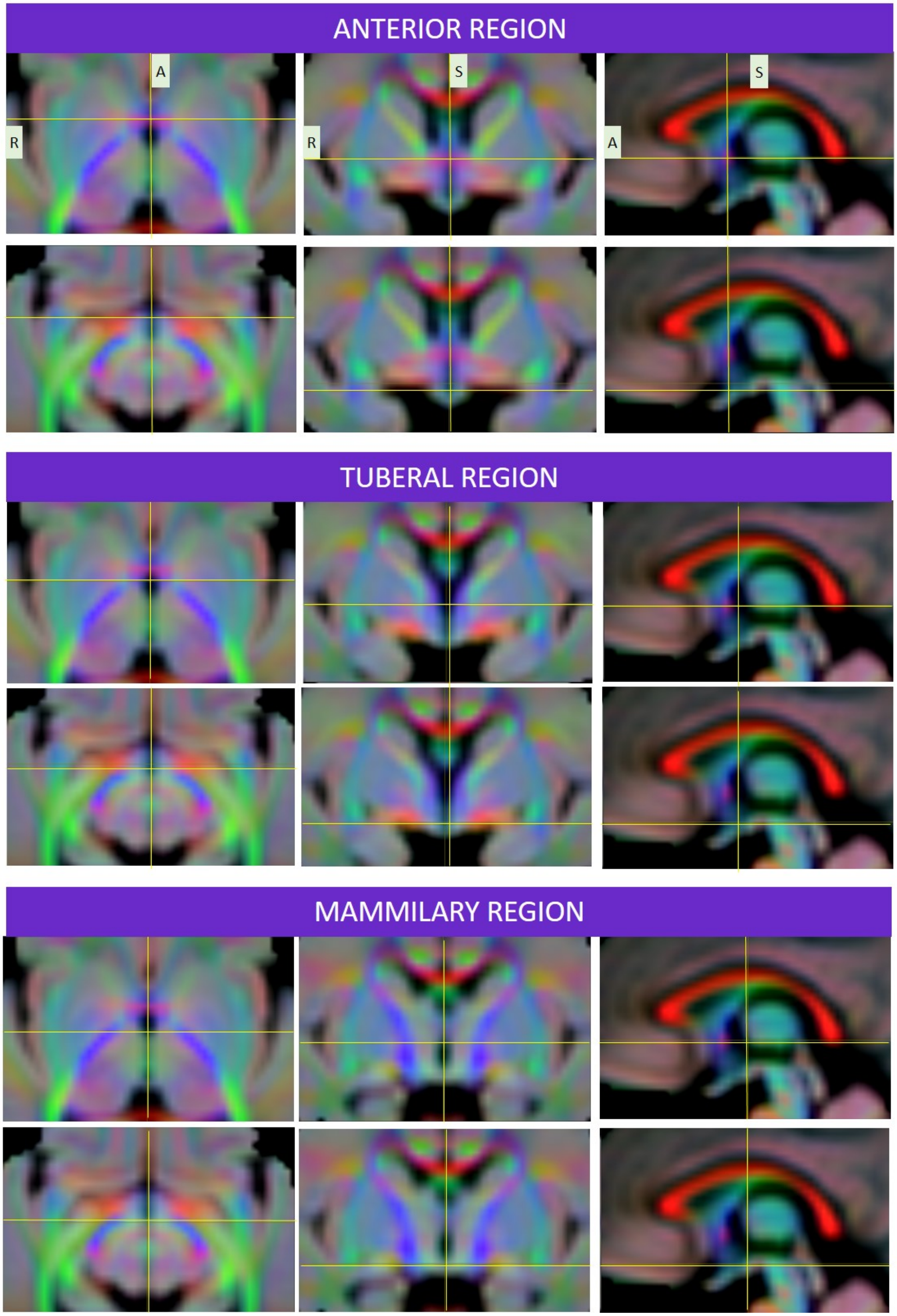


Figure S2: The FOD-derived directionally encoded color (DEC) map, weighted by the integral of the FOD was generated. This offers further visual information to place exclusion and inclusion gates for the tractography of the bundles of interest. The maps are presented in axial (first column), coronal (middle) and sagittal (right) views across the Anterior (Anterior commissure), Tuberal and Mammilary regions of the hypothalamus, with the top rows showing a superior aspect and the bottom row in each region the inferior aspect, as can be seen with the yellow cross-hairs. Red: left-right, blue:superior-inferior and green: anterior-posterior orientations.

Table S1: Tractography Protocols

| Tract | Seed | Seed direction | Inclusion | Exclusion | Cut-off | MaxMin length | Seed n | Other options |
|---|---|---|---|---|---|---|---|---|
| Am-Nacc | Amygdala | | Manual Nucleus Accumbens | | 0.10 | Max:45 | 5 million | |
| Cingulate | manual square mask in the cingulate gyrus | [0,1,0] | NA | NA | 0.10 | NA | 1 million | |
| DLF | manually drawn hypothalamus | | three manually drawn masks on medular, pontine tegmentum and midbrain periaqueductal gray in axial planes | manually drawn mask of fornix column on axial plane | 0.10 | Max:60 | 10 million | seed unidirectional |
| MFB | manually drawn VTA mask | | manually drawn NAcc mask | thalamus | 0.08 | Max:30 | 10 million | seed unidirectional |
| ST | manually drawn mask on the caudothalamic groove in the coronal plane similar to @Kamali2015 (assumed bed nucleus of the ST) | | amygdala | manually drawn mask of temporal stem in sagittal plane | 0.11 | | 20 million | |
| UF | manually drawn temporal stem of the UF [as in @Hau2016] using the DEC map to locate the ascending fibres | | medial and lateral temporal lobe, inferior and mid orbital frontal lobe | manually drawn hypothalamus, manually drawn mask of fornix column on coronal slice, thalamus | 0.10 | Min length: 20, Max 80 | 1 million | |
| vAMFP | manually drawn hypothalamus, | | amygdala | manually draw block over the fornix columns, manually drawn block in coronal plane to avoid streamlines to the ST | 0.13 | 30 | 5 million | seed unidirectional |

### 4 Protocols in probabilistic tractography

Since a few tracts had not been delineated in the neonate previously and others have used WM atlases based on manual delineation rather than tractography, certain tracts of interest (DLF, MFB) were first generated in a sample processed adult brain from the Human Connectome Project. This helped determine the location of regions of inclusion which have not yet been delineated in the neonate and also because of the resolution available in our cohort. We relied on structural anatomical atlases of the fetus and neonate (14) when possible.

Anatomical correspondence of the “core” body of a streamline bundle within each track was prioritised over the terminations of some fibre bundles which are known to “fan” out as they approach cortical surface (i.e. uncinate fasciculus) as we deemed this could influence the statistical metrics using mean FBA-derived measures. Further steps to minimise this was to utilise conservative fixel thresholding (count of streamline per fixel) within the fixel masks for each tract. The fixel masks are shown in Figure S3. Unless otherwise mentioned, all default software parameters were kept:

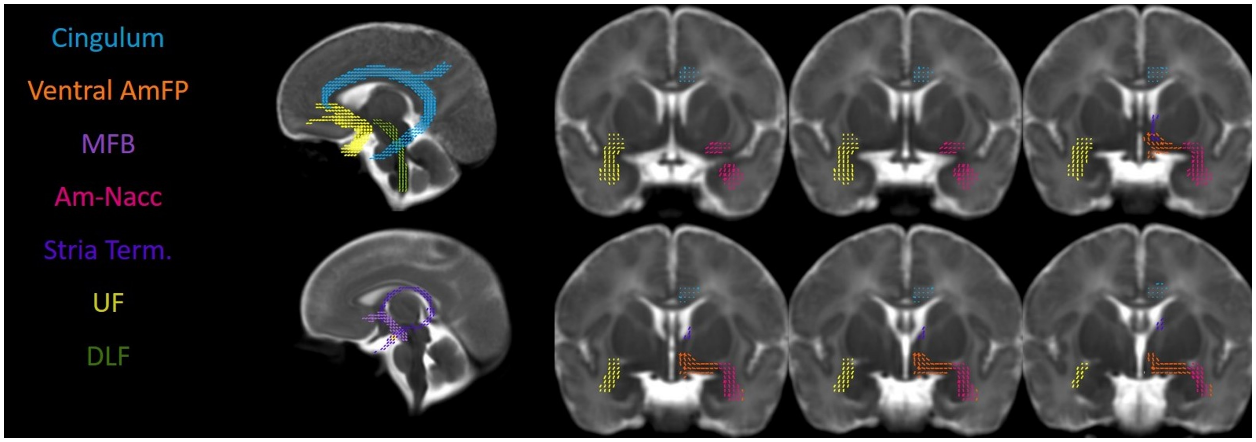


Figure S3: Fixel masks were generated and thresholded for each tract from which Fiber Density, Cross-section and Density X Cross-section were calculated. The coronal sections top left to bottom right: starts anterior to the anterior commissure to posterior.

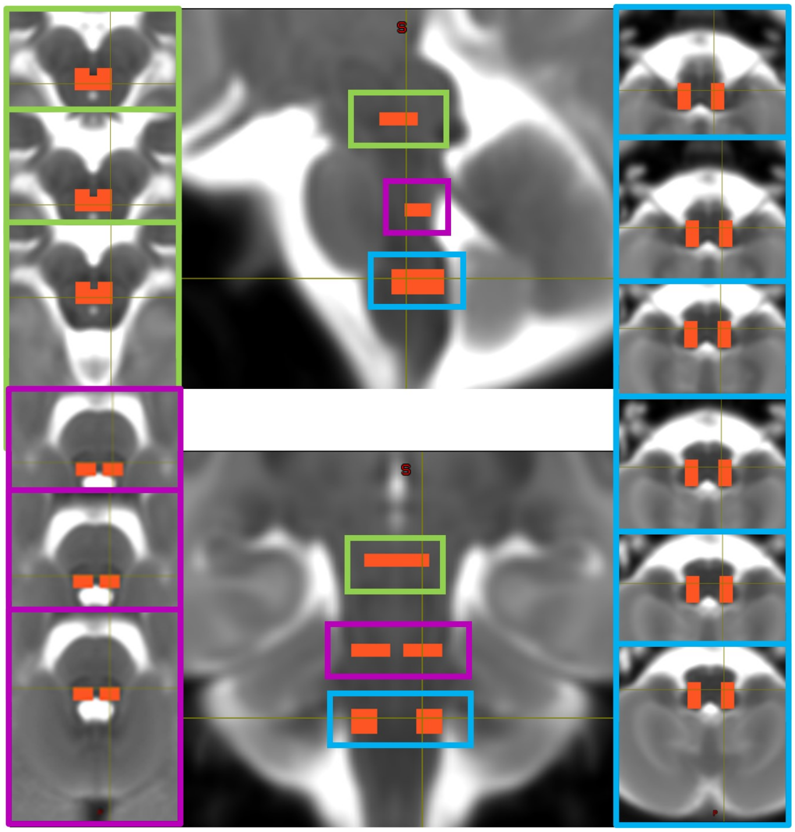

Figure S4: Regions of inclusion manually drawn for tractography of the Dorsal Longitudinal Fasciculus at the level of the midbrain periaqueductal grey matter (green), mid pons (fushia) and caudal medulla (blue) at a region assumed to include cranial nerve X (vagus) fibres. Central top image is in sagittal view and bottom in coronal view.

### 5 Normal-Weight pregnancies, birth and neonatal outcomes by sex

Table S2: Maternal and neonatal characteristics by sex in normal-weight pregnancies.

| | Male | Female | p |
|---|---|---|---|
| n | 77 | 60 | |
| **Mother** | | | |
| Mother's age (mean (SD)) | 33.78 (4.51) | 32.43 (5.57) | 0.120 |
| Mother's ethnicity (%) | | | 0.269 |
| White | 54 (72.0) | 37 (62.7) | |
| Black | 5 (6.7) | 5 (8.5) | |
| Asian | 6 (8.0) | 10 (16.9) | |
| Chinese | 6 (8.0) | 1 (1.7) | |
| Mixed | 1 (1.3) | 2 (3.4) | |
| Other | 3 (4.0) | 4 (6.8) | |
| Mother's first language is English (%) | 43 (56.6) | 31 (53.4) | 0.853 |
| Mother's age when last in FT education (median [IQR]) | 22.00 [21.00, 24.00] | 23.00 [21.00, 26.00] | 0.802 |
| Mother's BMI (median [IQR]) | 21.55 [20.23, 23.23] | 21.88 [20.17, 23.77] | 0.560 |
| Primiparous (%) | 47 (61.0) | 36 (60.0) | 1.000 |
| **Delivery** | | | |
| Meconium stained liquor (%) | 16 (21.3) | 16 (26.7) | 0.603 |
| Delivery Mode (%) | | | 0.988 |
| Elective caesarian section | 8 (10.4) | 6 (10.0) | |
| Emergency caesarian section - in labour | 18 (23.4) | 11 (18.3) | |
| Emergency caesarian section - not in labour | 3 (3.9) | 3 (5.0) | |
| Instrumental delivery - Forceps | 14 (18.2) | 12 (20.0) | |
| Instrumental delivery - Ventous | 6 (7.8) | 5 (8.3) | |
| Spontaneous vaginal delivery | 28 (36.4) | 23 (38.3) | |
| **Neonate** | | | |
| GA at birth weeks (mean (SD)) | 40.09 (1.01) | 40.46 (1.07) | 0.040 |
| Apgar 5 min (median [IQR]) | 10.00 [9.00, 10.00] | 10.00 [9.00, 10.00] | 0.283 |
| Baby head circumference (mean (SD)) | 34.81 (1.43) | 34.21 (1.63) | 0.022 |
| Baby birthweight (kg) (mean (SD)) | 3.43 (0.46) | 3.35 (0.49) | 0.374 |
| Birthweight centile (mean (SD)) | 51.76 (30.38) | 52.78 (29.92) | 0.844 |
| PMA at scan (mean (SD)) | 40.82 (1.33) | 41.12 (1.29) | 0.190 |

*Note:*
Mother's BMI (Body mass index) was booking BMI, GA: gestational age, PMA: Postmenstrual age (e.g. GA at birth + chronological postnatal age). Birthweight centile was calculated from INTERGROWTH-21st standards.

#### 5.1 Neonatal age distribution and path models

Path modeling relays other associations. As expected, TBV was predicted by sex (Male: 0.29[0.20 to 0.38]) and PMA at scan ([0.38[0.20 to 0.48]) but also birthweight centile (0.37[0.27 to 0.46]). Importantly, postnatal chronological age also associated positively with TBV (0.43[0.30 to 0.56]) even after accounting for the positive effect of PMA and male sex suggesting longer time outside the womb is also influences infant brain growth.

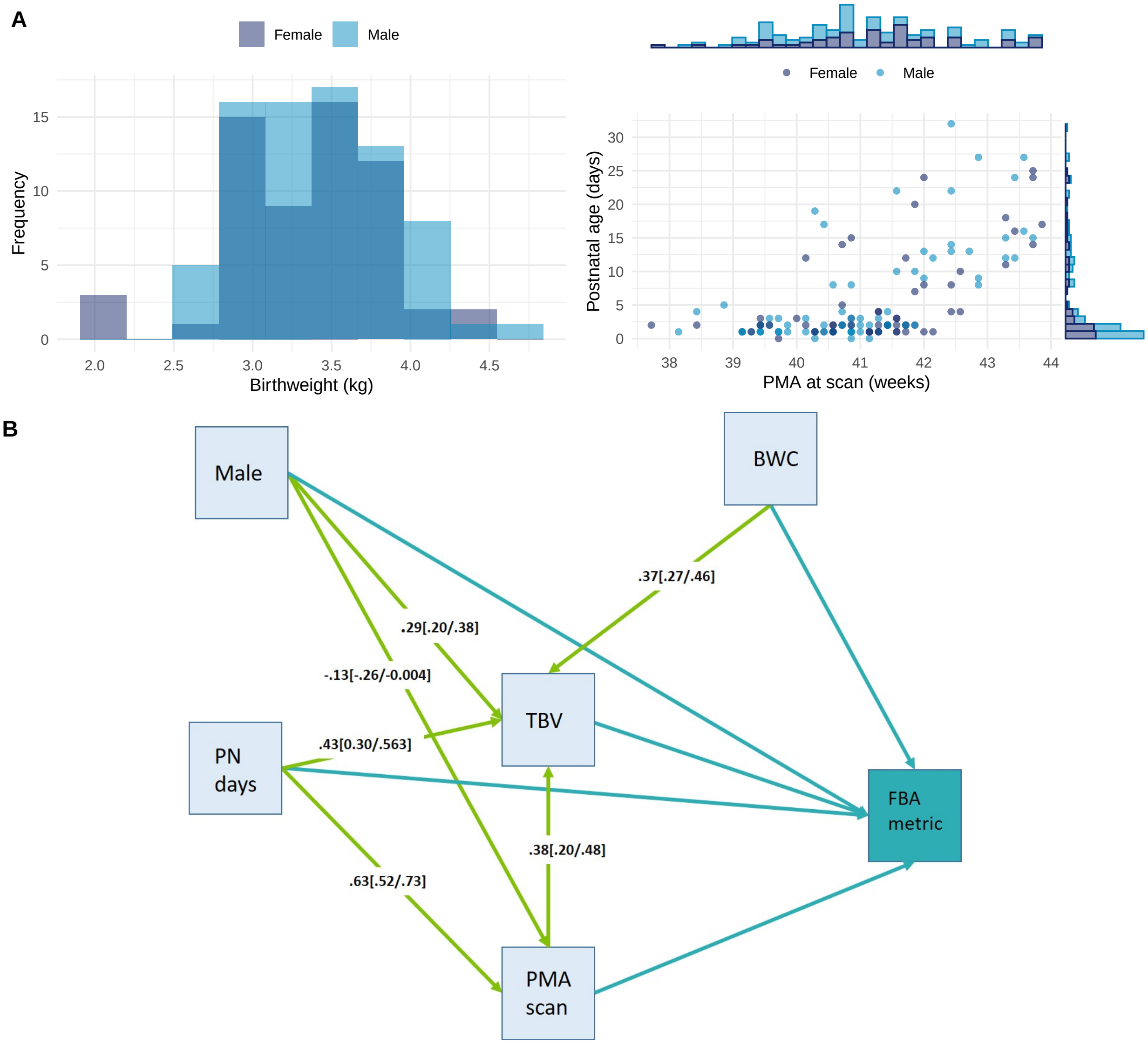


Figure S5: A: Neonatal characteristics in males (n=77) and females (n=60) born to normal-weight uncomplicated pregnancies. B: Path model from which regression beta coefficients [95%CIs] were obtained with direct paths (blue) and indirect paths (green) with Fixel-based metrics (fibre density, cross-section and density x cross-section). BWC: Birthweight Centile, FBA: Fixel-Based Analaysis, PMA: Postmenstrual age, PN: Postnatal, TBV: Total Brain Volume.

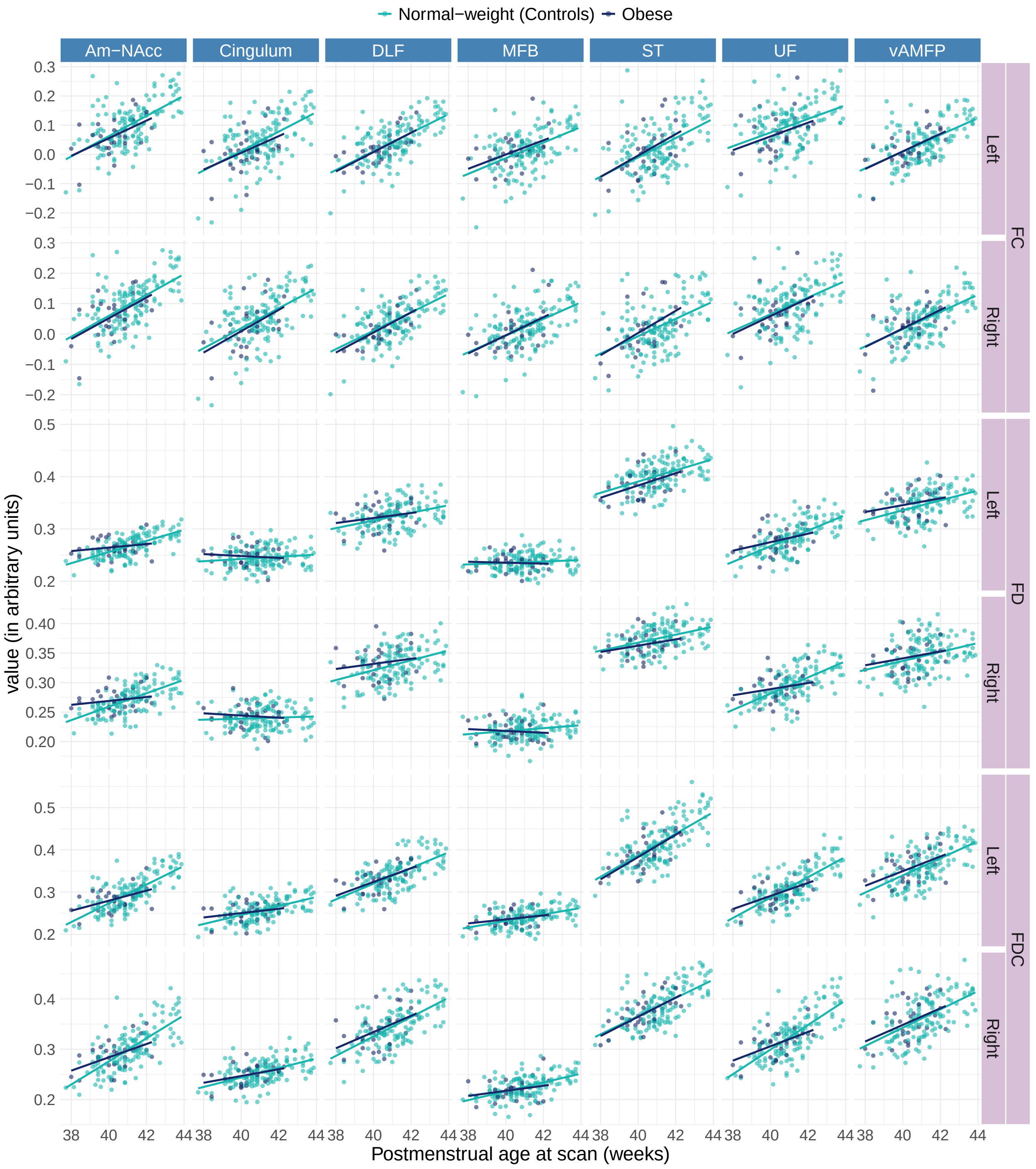


Figure S6: Correlations between fixel-based metrics and postmenstrual age at scan by exposure (normal-weight/obesity), sexes combined. Normal-weight: n=137 and obesity: n=28. FD:fibre density, FC:fibre cross-section, FDC:fibre density x fibre cross-section. DLF: Dorsal longitudinal fasciculus, MFB: Medial forebrain bundle, ST: Stria terminalis, UF: Unicinate fasciculus, vAMPF: Ventral amygdalofugal pathway.

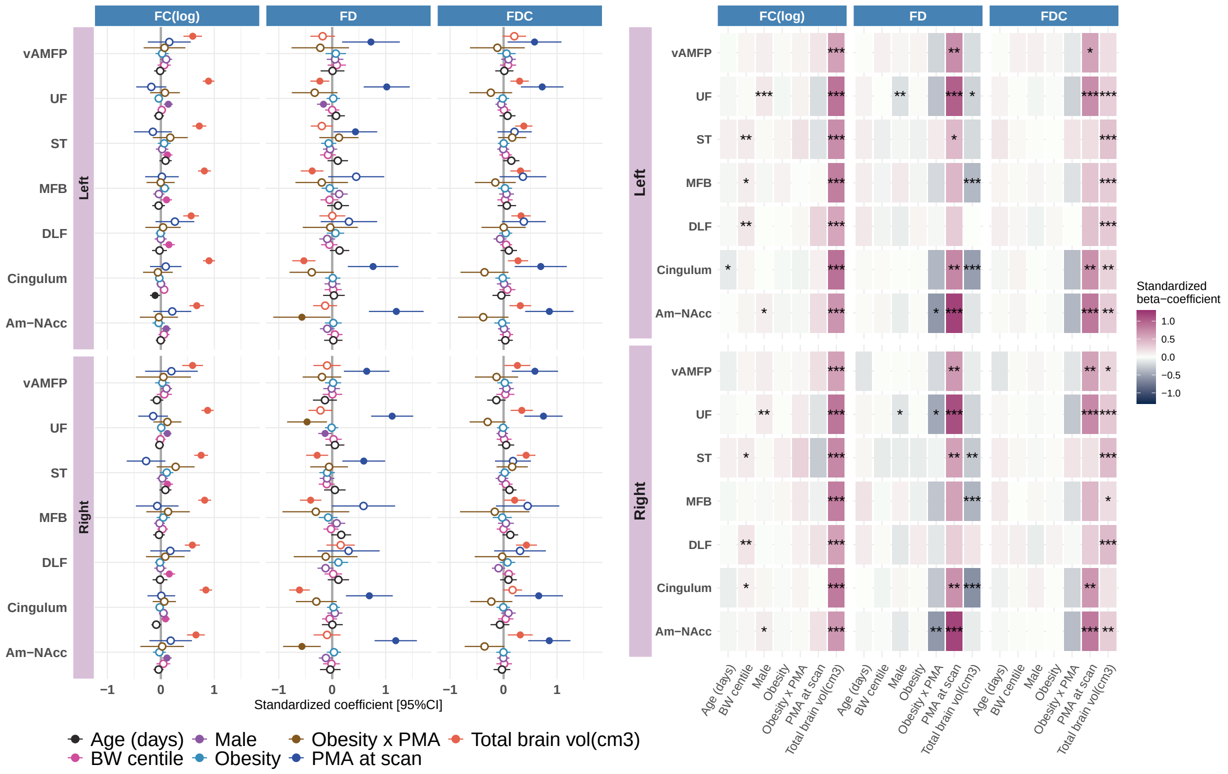


Figure S7: Figures represent the standardized beta coefficients obtained through path regression analyses with FD,FC and FDC as dependent variables. Left: circles are filled if the 95% confidence interval excludes 0. Right: stars denote significance levels *p<0.05,**p<0.01 and *** p<0.001. FD:fibre density, FC:fibre cross-section, FDC:fibre density x fibre cross-section. DLF: Dorsal longitudinal fasciculus, MFB: Medial forebrain bundle, ST: Stria terminalis, UF: Uncinate fasciculus, VAMPF: Ventral amygdalofugal pathway.

### 6 18-months outcomes

Table S3: Regression coefficients and p-values from path models on infant outcomes at 18 months.

| Predictor group | Predictor Tract | Hemi | Outcome Scale | Outcome Variable | std beta [95%CI] | p-value | FDR-corrected |
|---|---|---|---|---|---|---|---|
| tract | DLF | | Bayley | Receptive Lang | 0.28[0.13;0.43] | 0.00035 | 0.038 |
| Obesity | | | Bayley | Cognitive | -0.29[-0.45;-0.13] | 0.00046 | 0.038 |
| tract | Cingulum | | QCHAT | QCHAT total | 0.28[0.12;0.44] | 0.00073 | 0.040 |
| tract | DLF | | Bayley | Expressive Lang | 0.23[0.08;0.38] | 0.00213 | 0.088 |
| tract | Am-NAcc | | ECBQ | Surgency | 0.22[0.07;0.36] | 0.00402 | 0.132 |
| tract | Cingulum | | QCHAT | QCHAT total | 0.22[0.06;0.39] | 0.00795 | 0.199 |
| tract | Cingulum | | ECBQ | Negative Affect | 0.21[0.06;0.37] | 0.00844 | 0.199 |
| Obesity | | | Anthro | WAZ | 0.23[0.05;0.41] | 0.01207 | 0.249 |
| Obesity | | | Anthro | LHAZ | 0.2[0.02;0.38] | 0.01965 | 0.312 |
| tract | MFB | | ECBQ | Surgency | 0.19[0.03;0.36] | 0.02106 | 0.312 |
| tract | Am-NAcc | | ECBQ | Surgency | 0.19[0.03;0.34] | 0.02156 | 0.312 |
| tract | DLF | | Bayley | Expressive Lang | 0.18[0.03;0.33] | 0.02267 | 0.312 |
| tract | UF | | ECBQ | Surgency | 0.18[0.02;0.33] | 0.02760 | 0.327 |
| tract | DLF | | Bayley | Cognitive | 0.19[0.02;0.36] | 0.02774 | 0.327 |
| tract | vAMFP | | ECBQ | Negative Affect | 0.16[0.01;0.32] | 0.03827 | 0.401 |
| tract | UF | | ECBQ | Surgency | 0.16[0.01;0.32] | 0.03892 | 0.401 |
| tract | DLF | | QCHAT | QCHAT total | -0.19[-0.38;-0.01] | 0.04359 | 0.402 |
| tract | Cingulum | | Bayley | Receptive Lang | -0.18[-0.36;0] | 0.04380 | 0.402 |
| tract | Am-NAcc | | ECBQ | Effortful Control | -0.17[-0.34;0] | 0.04626 | 0.402 |
| tract | Cingulum | | Bayley | Cognitive | -0.15[-0.31;0] | 0.05386 | 0.444 |
| tract | DLF | | Bayley | Receptive Lang | 0.17[0;0.34] | 0.05743 | 0.451 |
| Obesity | | | ECBQ | Negative Affect | 0.18[-0.01;0.38] | 0.06922 | 0.516 |
| tract | UF | | ECBQ | Effortful Control | -0.17[-0.35;0.01] | 0.07186 | 0.516 |
| tract | DLF | | Bayley | Cognitive | 0.16[-0.02;0.33] | 0.07524 | 0.517 |
| tract | Am-NAcc | | ECBQ | Effortful Control | -0.14[-0.31;0.02] | 0.09433 | 0.623 |
| tract | Cingulum | | Bayley | Receptive Lang | -0.15[-0.33;0.03] | 0.10696 | 0.632 |
| tract | MFB | | Anthro | LHAZ | -0.14[-0.31;0.03] | 0.11297 | 0.632 |
| tract | Cingulum | | Bayley | Cognitive | -0.12[-0.27;0.03] | 0.11572 | 0.632 |
| tract | Cingulum | | Bayley | Expressive Lang | -0.15[-0.34;0.04] | 0.11923 | 0.632 |
| tract | UF | | ECBQ | Effortful Control | -0.14[-0.32;0.04] | 0.12235 | 0.632 |
| tract | Am-NAcc | | Bayley | Expressive Lang | 0.13[-0.04;0.3] | 0.12577 | 0.632 |
| tract | ST | | ECBQ | Effortful Control | -0.12[-0.27;0.03] | 0.12626 | 0.632 |
| tract | Am-NAcc | | CBCL | Internalising | -0.12[-0.27;0.04] | 0.12876 | 0.632 |
| tract | ST | | ECBQ | Effortful Control | -0.12[-0.26;0.03] | 0.13028 | 0.632 |
| tract | Cingulum | | Bayley | Expressive Lang | -0.13[-0.31;0.04] | 0.14048 | 0.662 |
| tract | MFB | | Anthro | WAZ | -0.11[-0.27;0.04] | 0.15474 | 0.694 |
| tract | UF | | ECBQ | Negative Affect | 0.1[-0.04;0.24] | 0.15795 | 0.694 |
| tract | Cingulum | | ECBQ | Negative Affect | 0.11[-0.04;0.27] | 0.16092 | 0.694 |
| tract | DLF | | QCHAT | QCHAT total | -0.13[-0.32;0.05] | 0.16392 | 0.694 |
| tract | MFB | | Bayley | Expressive Lang | -0.12[-0.28;0.05] | 0.17830 | 0.735 |
| Obesity | | | Bayley | Receptive Lang | -0.12[-0.3;0.06] | 0.18750 | 0.755 |
| tract | Cingulum | | Anthro | WAZ | -0.11[-0.29;0.06] | 0.20337 | 0.799 |
| tract | ST | | QCHAT | QCHAT total | 0.1[-0.06;0.26] | 0.20984 | 0.805 |
| tract | MFB | | CBCL | Internalising | -0.1[-0.26;0.06] | 0.23053 | 0.843 |
| Obesity | | | ECBQ | Effortful Control | 0.09[-0.06;0.24] | 0.23960 | 0.843 |
| tract | vAMFP | | QCHAT | QCHAT total | 0.1[-0.07;0.27] | 0.24281 | 0.843 |
| tract | MFB | | CBCL | Externalising | -0.09[-0.25;0.06] | 0.24850 | 0.843 |
| tract | Am-NAcc | | CBCL | Externalising | 0.1[-0.07;0.27] | 0.25179 | 0.843 |
| tract | DLF | | CBCL | Internalising | -0.12[-0.32;0.08] | 0.25179 | 0.843 |
| tract | vAMFP | | ECBQ | Effortful Control | -0.11[-0.3;0.08] | 0.25554 | 0.843 |

[a] FDR corrected at p<0.05 on a total of 165 regression paths. Only top 50 shown.
